# FOLR1-targeted Actinium-225-based Alpha-particle Therapy Eliminates Ovarian Cancer

**DOI:** 10.1101/2025.10.27.684721

**Authors:** Neetu Singh, Esther Need, Ayden Berndt, Matthew Goff, Lydia Wilson, Firas Mourtada, Feng Guo, Tara Mastren, Taslim Al-Hilal, Anil K Sood, Amit Maity, Scott C Miller, Satoshi Minoshima, Shreya Goel, Sixiang Shi

## Abstract

Despite the advancement in therapies, ovarian cancer treatment is challenging due to poor prognosis and high relapse associated with acquired resistance. Targeting overexpression of FOLR1 in ovarian cancers has proven to be an attractive strategy. The recent FDA approval of FOLR1 targeted antibody drug conjugate has shown promising results albeit resistance with repeated use appears inevitable. Emerging targeted alpha-particle therapies, particularly Actinium-225 (^225^Ac), for treating refractory cancers have opened avenues for improved therapeutic options. The success of alpha-particle therapy relies on tumor specific delivery of the alpha emitters. Herein we describe the first example of FOLR1-targeted ^225^Ac alpha-particle therapy for treatment of ovarian cancer. Longitudinal PET imaging demonstrated high tumor-specific uptake of αFOLR1 in SKOV3 xenografts. FOLR1-targeted ^225^Ac demonstrated high therapeutic efficacy achieving marked tumor regression, 80% survival and 40% complete response. The therapy resulted in tumor specific double stranded DNA damage, and no obvious toxicity was observed in normal tissues. Estimated human dosimetry showed high absorbed dose for tumor and minimal absorbed dose for healthy tissues establishing its safety. In totality, FOLR1-targeted ^225^Ac alpha-particle therapy is an efficacious and safe treatment with high feasibility for clinical translation to fight against ovarian cancer.

## Main Text

Ovarian cancer is the fifth leading cause of cancer associated death among women and often presented at advanced stages with poor prognosis and limited therapeutic treatment options^1,2^. Patients with ovarian cancer experience high morbidity and mortality with median survival rates of 40-50% at stage III and only ∼17% at stage IV ^3,4^. Despite the advances in surgical interventions and well-designed treatment regimens with chemotherapy agents, most patients relapse within two years and many recurrent tumors are taxane and platinum resistant^4^. Intrinsic or acquired resistance to therapies limits their benefit in advanced ovarian cancers, highlighting the critical unmet need to explore novel therapeutic directions^5^.

Targeted alpha-particle therapy (TAT) has emerged as a promising approach with demonstrable clinical success, especially for the cancer types that cannot be completely removed by surgical resection and are resistant to frontline chemo- and immuno-therapies. Compared with beta emitters (which primarily induce single-stranded DNA breaks in target cells), the alpha emitter, Actinium-225 (^225^Ac), has higher linear energy transfer. This results in more lethal double-stranded DNA breaks that are difficult to repair and can effectively kill tumor cells. As such, TAT remains effective despite conventional cellular resistance mechanisms^6,7^. Additionally, ^225^Ac has a short path length (< 100 µm) in tissues, which confines its impact to neighboring cells and significantly reduces the off-target toxicity^8-10^. Therefore, ^225^Ac-based TAT is a promising approach for treating cancers, especially those that are late-stage and therapy-resistant. On the other hand, high linear energy transfer and short path length of ^225^Ac necessitate accurate delivery to tumor cells, as off-target delivery will lead to unwanted side effects and compromised efficacy. Thus, success of TATs is heavily dependent on selection of optimized molecular targets that possess excellent tumor specificity and optimal pharmacokinetic profiles *in vivo*. Although a number of TAT approaches are currently under preclinical and clinical investigation for multiple cancer types, it has not been applied to the treatment of ovarian cancer^11^.

Folate receptor 1 (encoded by FOLR1 gene) or Folate receptor α is an extensively studied molecular target for ovarian cancer therapy development^12,13^. FOLR1 is a cell-surface protein that binds to folic acid and its derivatives, which plays a key role in metabolic processes, DNA replication, and cell division. FOLR1 exhibits elevated expression in 70% of primary and 80% of recurrent ovarian cancer with minimal expression in non-malignant tissues^14^. FOLR1 can internalize relatively large molecules rendering it suitable for development of targeted therapies^12,15^. Recently, the FDA approved the first FOLR1 targeted antibody-drug conjugate, Mirvetuximab Soravtansine (a tubulin inhibitor), for the treatment of platinum-resistant ovarian cancers^16^. Mirvetuximab Soravtansine demonstrated a high response rate (34% vs 16% in standard chemotherapy) in platinum resistant ovarian cancer patients and improved overall survival by 4 months^17^. Inspired by the pioneering success of Mirvetuximab Soravtansine in the clinic, we hypothesized that FOLR1 can be a promising target for developing next-generation radiopharmaceuticals against ovarian cancers.

Herein, we combine the tumor specificity offered by FOLR1-targeted antibody (αFOLR1) and potent antitumor effects of ^225^Ac to develop a novel FOLR1-targeted ^225^Ac-based TAT for ovarian cancers. In this study, we present *in vitro* and *in vivo* characterization of FOLR1 targeted ^225^Ac-Macropa-αFOLR1 in the SKOV3 ovarian cancer model. Our data demonstrate exceptional *in vivo* pharmacokinetic profile and tumor specificity of αFOLR1 for ^225^Ac delivery. ^225^Ac-Macropa-αFOLR1 demonstrated remarkable antitumor effects, survival rate and safety compared to folate-targeted antibody-chemo drug conjugates. Taken together, we report the first successful example of FOLR1-targeted ^225^Ac-based alpha therapy for ovarian cancer as an effective and well tolerated treatment strategy with high clinical relevance.

## Results

### Transcriptome analysis confirms FOLR1 overexpression in ovarian cancer

By analyzing the TCGA database, we confirmed that ovarian cancer (OV) has the highest FOLR1 expression across all cancer types (**Figure 1A**). Comparable results about significant overexpression of FOLR1 in ovarian/fallopian cancer can also be found in Cancer Cell Line Encyclopedia (CCLE) datasets (**Supplementary Figure 1**). Radar plot visualization (**Figure 1B**) further emphasized that FOLR1 expression in OV exceeds all other tumor types as well as the paired adjacent normal tissues, consistent with the aforementioned findings. Importantly, the FOLR1 expression in ovarian carcinoma was found to be significantly elevated in comparison to normal ovary tissues, derived from The Adult Genotype Tissue Expression (GTEx) Project database (**Figure 1C**).

**Figure 1.**
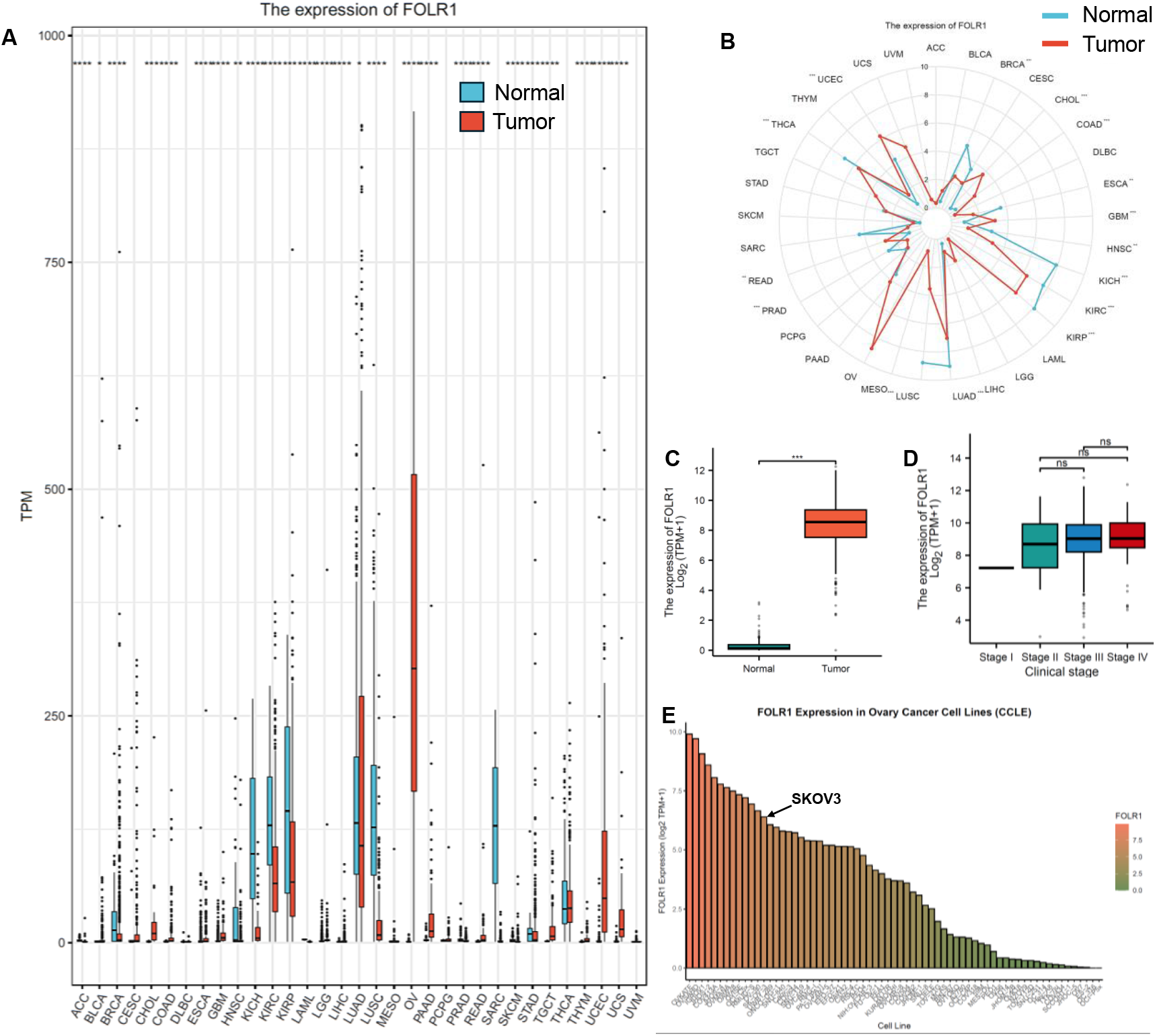
Transcriptome analysis of FOLR1 expression. (A) Transcripts per million (TPM) indicating FOLR1 expression across different cancer tissues from TCGA database and normal tissues from GTex database. Some cancer types (for example, Sarcoma (SARC), Skin cutaneous melanoma (SKCM), Thymoma (THYM), Adrenocortical carcinoma (ACC), Lymphoid Neoplasm Diffuse Large B-cell Lymphoma (DLBC), Acute Myeloid Leukemia (LAML), Brain lower grade glioma (LGG), Mesothelioma (MESO)) had insufficient normal controls (n < 3 or SD = 0) and were excluded from statistical comparisons though included in visualizations. (B) Radar plot showing the highest FOLR1 expression in ovarian tumors (C) Comparison of ovarian tumor and normal ovary indicating significant high expression in tumor tissues (D) Expression of FOLR1 at different clinical stages showing no significant changes in expression levels. (E) FOLR1 expression levels in different ovarian cancer cell lines evaluated from CCLE database.

Interestingly, we found that FOLR1 expression is homogenously elevated in FOLR1-postive ovarian cancer at all clinical stages (Wilcoxon rank-sum test) (**Figure 1D**). Detailed correlation analyses (data not shown) between FOLR1 and other clinical features, including tumor status, primary therapy outcome, histologic grade, lymphovascular invasion, and survival endpoints (overall survival (OS), disease specific survival (DSS), progression free interval (PFI)) did not reveal any strong associations. Across multiple survival analyses in TCGA-OV (which includes only cystadenocarcinoma), FOLR1 expression was not associated with prognosis. Dichotomization (high vs. low), tertile, and quartile-based stratification consistently showed no significant differences in OS, DSS, or PFI. These observations indicate that as-reported FOLR1-based TAT is applicable and suitable for all FOLR1-expressing ovarian cancer at all stages, highlighting its strong relevance to broad clinical translation and value for commercialization.

In addition to tumor tissues, we analyzed the FOLR1 transcriptome in ovarian cancer cell lines. We observed heterogeneity with several cell lines showing high to moderate levels of FOLR1 expression (**Figure 1E**). In our study, we selected the SKOV3 ovarian cancer cell line that presents moderate FOLR1 expression. We reasoned that if FOLR1-based TAT is effective in SKOV3 model, it should be also effective in other tumor models with higher FOLR1 expression.

### FOLR1 is overexpressed on SKOV3 cells and tumor tissues

The expression of FOLR1 on SKOV3 ovarian cancer cell line was evaluated by flow cytometry. SKOV3 cells incubated with Alexafluor 488-labeled anti-FOLR1 antibody (αFOLR1) demonstrated a 4-fold increase in median fluorescence intensity (**Figure 2A, B**) compared to Alexafluor 488 labeled isotype control, indicating overexpression of FOLR1 on SKOV3 cells. We further evaluated FOLR1 expression in SKOV3 subcutaneous ovarian cancer xenografts by immunofluorescence (**Figure 2C**). The tumor tissues stained with Cy5-αFOLR1 exhibited elevated signal intensity (indicated by red) suggesting high FOLR1 expression compared to Cy5-Isotype. Notably, normal ovary tissues did not show any marked enhancement in signal. Furthermore, FOLR1 expression on liver, spleen and kidney was minimal. Our results indicated SKOV3 ovarian cancers have high FOLR1 expression and FOLR1 is a suitable biomarker for *in vivo* targeting of SKOV3 ovarian cancers.

**Figure 2.**
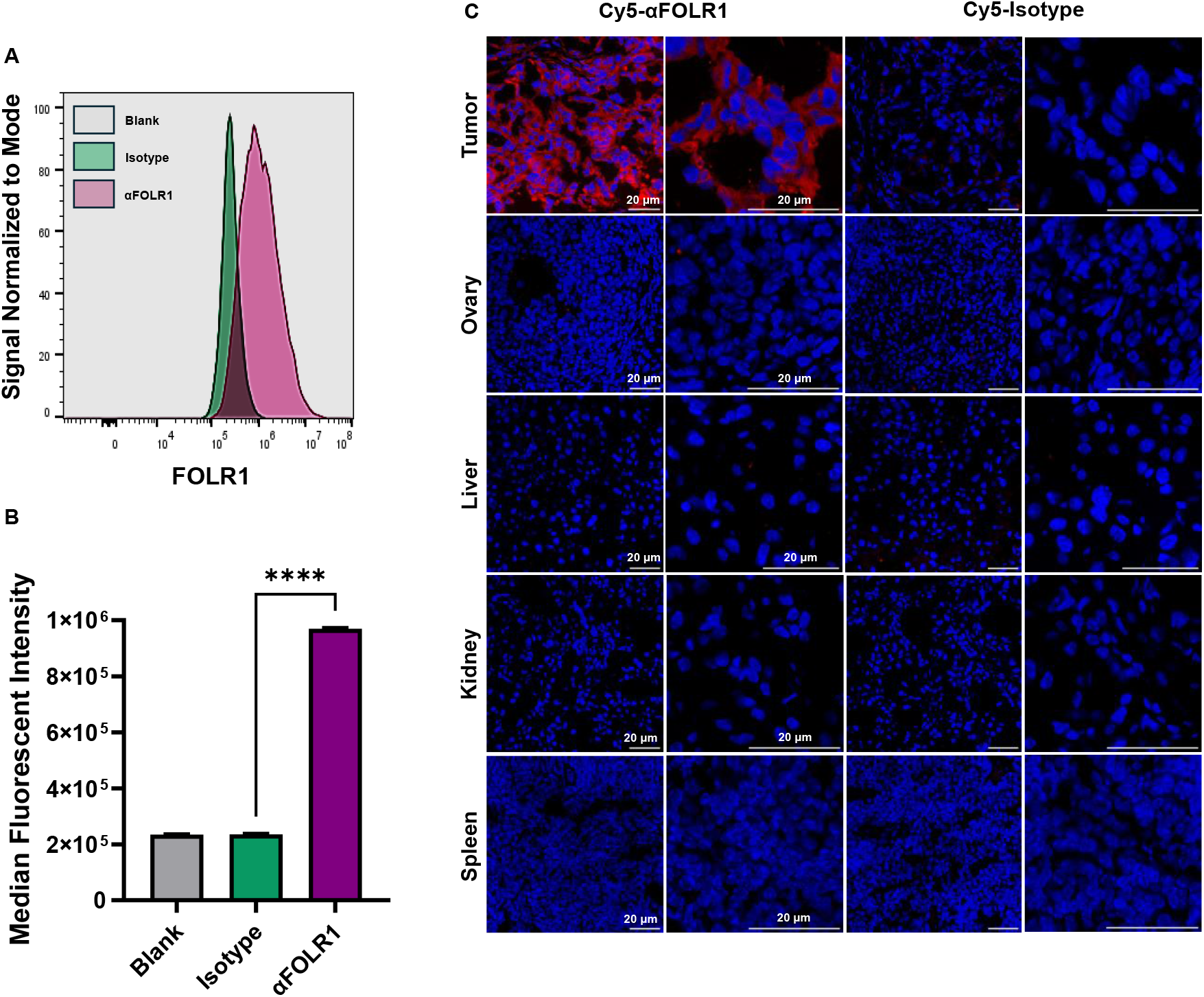
Assessment of FOLR1 expression in SKOV3 ovarian cancer model. (A, B) FOLR1 expression in SKOV3 ovarian cancer cell in vitro by flow cytometry. (C) Expression of FOLR1 in SKOV3 xenograft tissues evaluated by immunofluorescence. Statistical analysis was performed using unpaired t-test (n = 3) *p < 0.05, ** p < 0.01, ***p < 0.001, ****p < 0.0001, ns = non-significant (p > 0.05).

### αFOLR1 exhibited excellent tumor uptake and attractive pharmacokinetics *in vivo*

The success of TAT is heavily dependent on the design of optimal targeting ligands that possess excellent molecular specificity and optimal pharmacokinetic profiles *in vivo*. To ensure the success of FOLR1-based TAT, we employed longitudinal positron emission tomography (PET) to examine the *in vivo* tumor uptake and biodistribution of αFOLR1 antibody. In brief, αFOLR1 antibody and corresponding isotype IgG (Isotype) were radiolabeled with Zirconium-89 (^89^Zr) using deferoxamine (DFO) as a chelator using a standard radiolabeling approach (**Figure 3A**)^18^. The reaction was carried out at pH 7 and 37°C for a duration of 2h. The kinetic labeling (**Figure 3B**) yield was assessed throughout the reaction. High labeling yield was achieved within 1h incubation for αFOLR1 (99.2 ± 0.14 %) as well as the Isotype control (96.24 ± 0.37%). *In vitro* stability of ^89^Zr-DFO-labeled antibodies was evaluated in mouse serum and saline buffers (pH 7.4) at room temperature. As shown in (**Figure 3C, D**) both ^89^Zr-DFO-αFOLR1 and ^89^Zr-DFO-Isotype remained stable for 72 h in serum (>98%) and saline (>95%).

**Figure 3.**
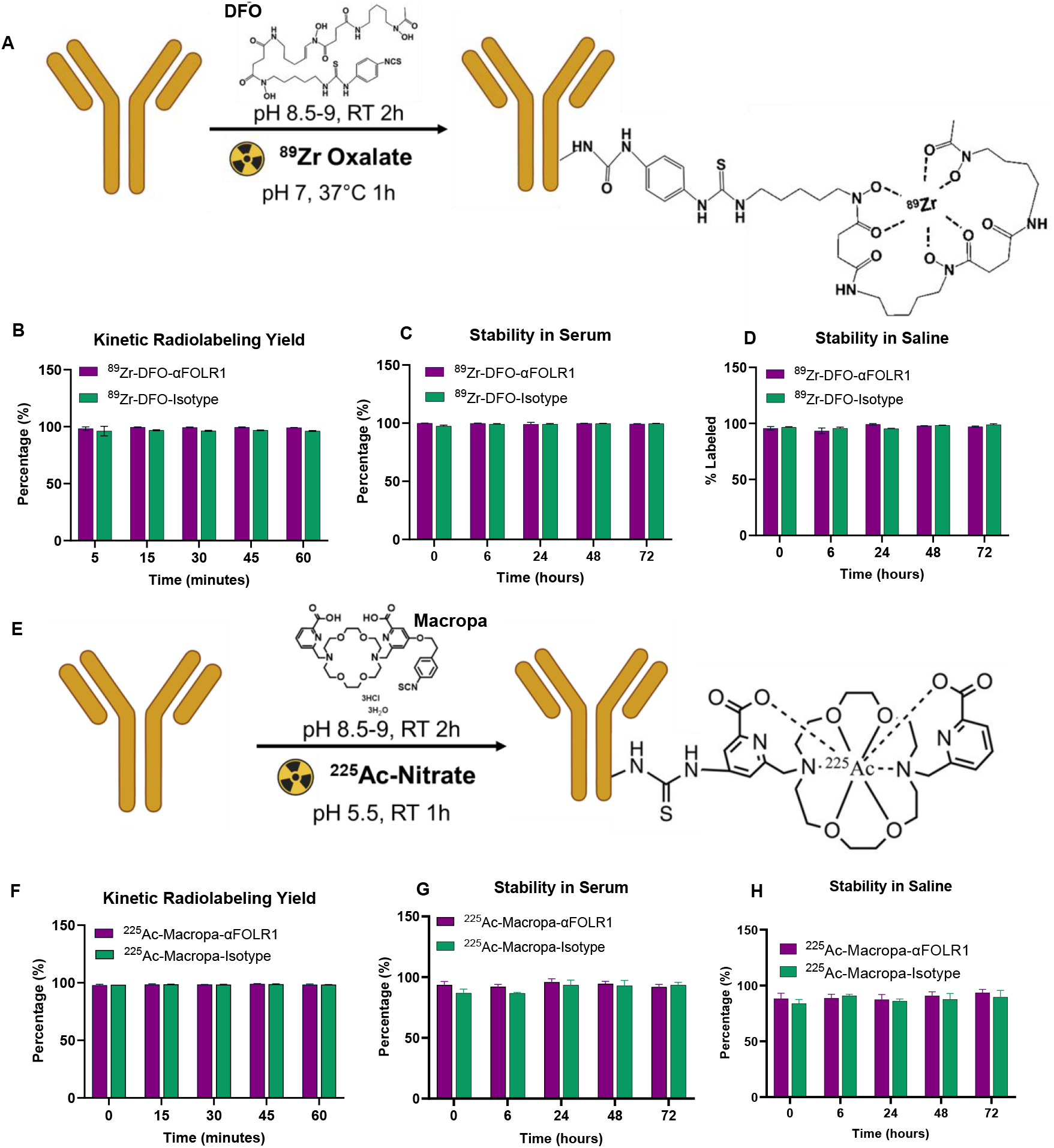
Development and Characterization of FOLR1-targeted Radiotracers. (A) Schematic for radiolabeling of αFOLR1 with Zirconium-89. Evaluation of ^89^Zr-DFO-αFOLR1 for (B) Kinetic radiolabeling yield, (C) radiolabeling stability in serum (D) radiolabeling stability in saline. (E) Schematic for development of ^225^Ac-Macropa-αFOLR1 radioconjugate. Assessment of (F) kinetic radiolabeling yield (G) radiolabeling stability in serum (H) radiolabeling stability in saline of ^225^Ac-Macropa-αFOLR1.

Serial non-invasive PET scans were used for visual and quantitative assessment of whole-body biodistribution and pharmacokinetics of ^89^Zr-DFO-αFOLR1. The corresponding isotype IgG (^89^Zr-DFO-Isotype) was used as a control to understand the influence of nonspecific tumor uptake via enhanced perfusion and retention in tumor; i.e., any tumor uptake greater than the isotype control can be attributed to molecular specificity for FOLR1 receptor. The longitudinal maximum intensity projection (MIP) images (**Figure 4A**) demonstrated an increasing uptake of ^89^Zr-DFO-αFOLR1 at the tumor site over 72 h post injection (p.i.). The time activity curves (**Figure 4B**) indicated ^89^Zr-DFO-αFOLR1 achieved significantly higher tumor uptake (39.17± 5.26 %ID/g) compared to ^89^Zr-DFO-Isotype (11.2 ± 1.78 %ID/g) at 72 h p.i. ^89^Zr-DFO-αFOLR1 uptake was minimal in the liver (7.2 ± 3.6 %ID/g) and spleen (6.4 ± 2.8 %ID/g). At 72 h p.i., *ex vivo* biodistribution (**Figure 4C**) of activity assessed by gamma counting corroborated the *in vivo* findings with both ^89^Zr-DFO-αFOLR1 (48.9 ± 5.5 %ID/g) and ^89^Zr-DFO-Isotype (10.09 ± 1.2 %ID/g), validating the accuracy of PET imaging. Moreover, the uptake ^89^Zr-DFO-αFOLR1 in liver and spleen was significantly less than ^89^Zr-DFO-Isotype control, indicating high FOLR1 specificity and potential low toxicity of FOLR1-based TAT.

**Figure 4.**
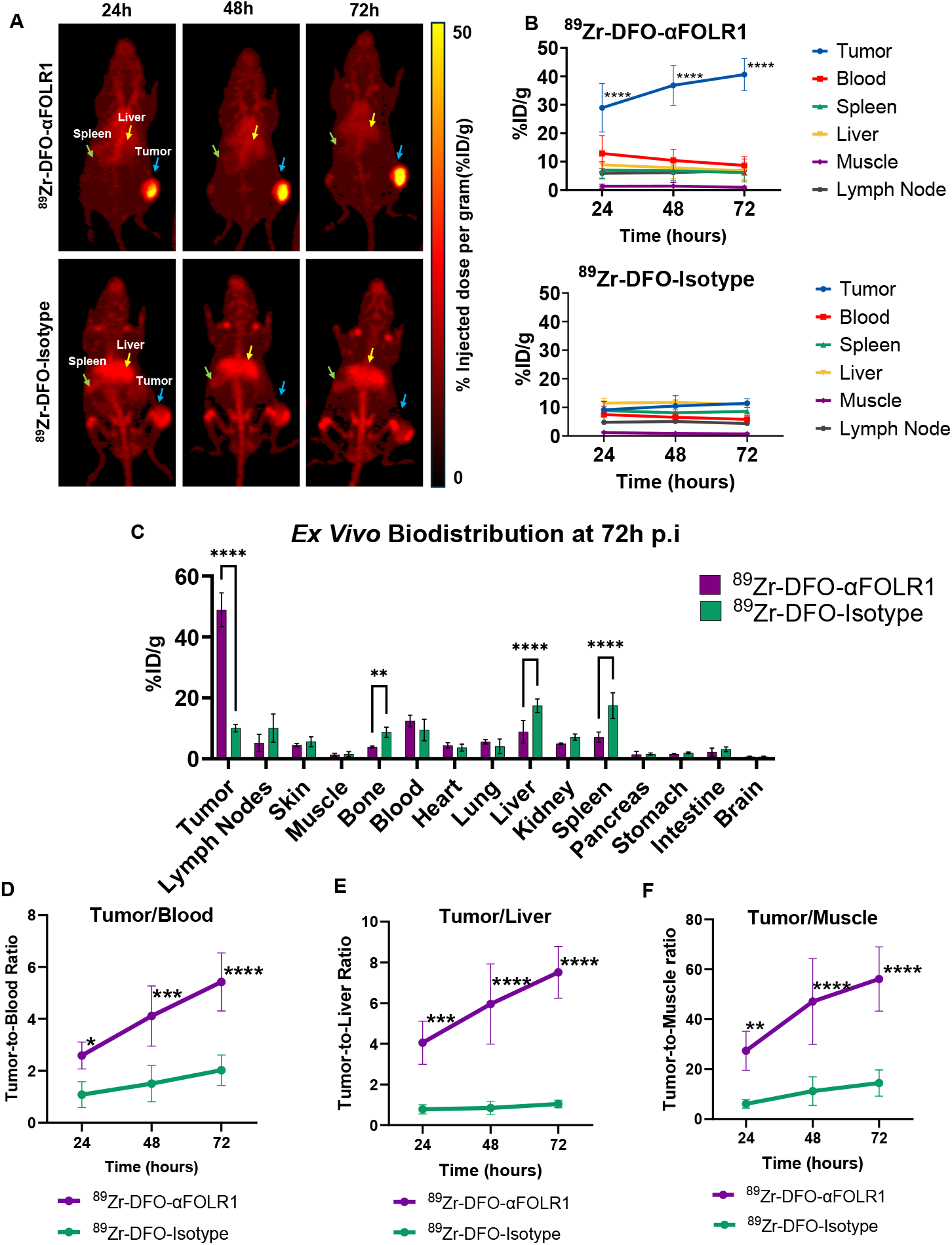
In vivo PET Images, Pharmacokinetics and Biodistribution of αFOLR1. (A) Longitudinal MIP PET images demonstrating increasing tumor uptake of ^89^Zr-DFO-αFOLR1 (upper panel) compared to ^89^Zr-DFO-Isotype (lower panel). (B) Time activity curves generated by region-of-interest analysis of PET images for ^89^Zr-DFO-αFOLR1 (upper panel) and ^89^Zr-DFO-Isotype (lower panel). (C) Ex vivo biodistribution profile at 72 h p.i. Tumor to (D) Blood (E) Liver and (F) Muscle signal ratio for ^89^Zr-DFO-αFOLR1 and ^89^Zr-DFO-Isotype. Statistical analysis was performed using unpaired t-test and two-way ANOVA (multiple comparison) (n = 4) *p < 0.05, ** p < 0.01, ***p < 0.001, ****p < 0.0001, ns = non-significant (p > 0.05).

Additionally, we evaluated the tumor tissue to background organ signal ratio to further delineate the targeting specificity. Our results indicated significantly high tumor to blood ratio (**Figure 4D**) attained by ^89^Zr-DFO-αFOLR1 (5.4 ± 1.2) in comparison to ^89^Zr-DFO-Isotype (2.02 ± 0.58) at 72 h p.i., further providing evidence for molecular targeted tumor accumulation of our construct. Of note, ^89^Zr-DFO-αFOLR1 demonstrated an increasing trend in tumor accumulation and remained in circulation at 72 h p.i. (∼ 10%ID/g in blood); thus, we expect that this ratio will further increase at later timepoints, as more antibody clears from the blood and accumulates in tumors. By contrast, the isotype kinetics plateaued over this period. We also observed high tumor to liver ratio (**Figure 4E**) in ^89^Zr-DFO-αFOLR1 (7.5 ± 1.2) treated group when compared to ^89^Zr-DFO-Isotype (1.03 ± 0.1). In addition, tumor to muscle ratio of ^89^Zr-DFO-αFOLR1 (56.13 ± 12) treated group was significantly higher compared to ^89^Zr-DFO-Isotype (14.45 ± 5.22) (**Figure 4F**).

### ^225^Ac-Macropa-αFOLR1 exhibits robust therapeutic efficacy in SKOV3 ovarian cancer xenografts

Robust and stable ^225^Ac labeling has been a significant obstacle in developing ^225^Ac-based TAT. Conventional DOTA ((1,4,7,10-Tetraazacyclododecane-1,4,7,10-tetrayl tetraacetic acid)-based chelation requires high temperature and results in low labeling yields, which significantly limits its potential application in antibody-based TAT^19^. To overcome these challenges, αFOLR1 was conjugated to an N-Hydroxysuccinimide-modified bifunctional chelator Macropa 6-((16-((6-carboxypyridin-2-yl)methyl)-1,4,10,13-tetraoxa-7,16-diazacyclooctadecan-7-yl)methyl)-4-(4-isothiocyanatophenethoxy)picolinic acid (Ratio Therapeutics, Boston, MA) to allow room-temperature and high-yield radiolabeling of ^225^Ac (**Figure 3E**). Macropa-αFOLR1 was purified and stored in ammonium acetate buffer (pH 7) until radiolabeling. The conjugate was labeled with ^225^Ac with a high yield of > 95% (**Figure 3F**). The radiochemical purity of the final construct was > 98% as determined by instant thin layer chromatography. ^225^Ac-Macropa-αFOLR1 demonstrated excellent stability in serum (> 92 %) (**Figure 3G**) and saline (> 88%) (**Figure 3H**) for up to 72 h, providing enough time for the stable radioconjugate to accumulate at tumor site. Taking together, the successful, robust, and stable room-temperature labeling of ^225^Ac is critical to ensure successful targeted delivery of ^225^Ac to the tumor *in vivo*, without pre-target detachment or modification of antibody structure. Our results highlight the readiness and strong potential of as-designed constructions for clinical translation.

We next assessed the therapeutic efficacy ^225^Ac-Macropa-αFOLR1 in subcutaneous SKOV3 tumor bearing mice (**Figure 5A**). Mice were treated intravenously with a single high dose (1 µCi) of ^225^Ac-Macropa-αFOLR1 or ^225^Ac-Macropa-Isotype. A group of mice that received no treatment served as control. Our results demonstrated no significant change in relative tumor volume (**Supplementary Figure 2A-C**) and body weight (**Supplementary Figure 2D-F**) between the different treatment groups. *Ex vivo* biodistribution assessments at Day 14 (**Supplementary Figure 3**) post treatment showed a 30% tumor uptake of ^225^Ac-Macropa-αFOLR1 compared to 10% uptake of ^225^Ac-Macropa-Isotype, underscoring high targetability and tumor specificity. However, we observed premature animal morbidity in both ^225^Ac-Macropa-αFOLR1, and ^225^Ac-Macropa-Isotype treated groups around 10 days after treatment, potentially due to radiotoxicity and poor tolerance. The results of our first trial with single high dose (1 µCi) suggest that a better low-dose therapeutic regimen should be developed to reduce the radiotoxicity without compromising tumor deposition of the isotope.

**Figure 5.**
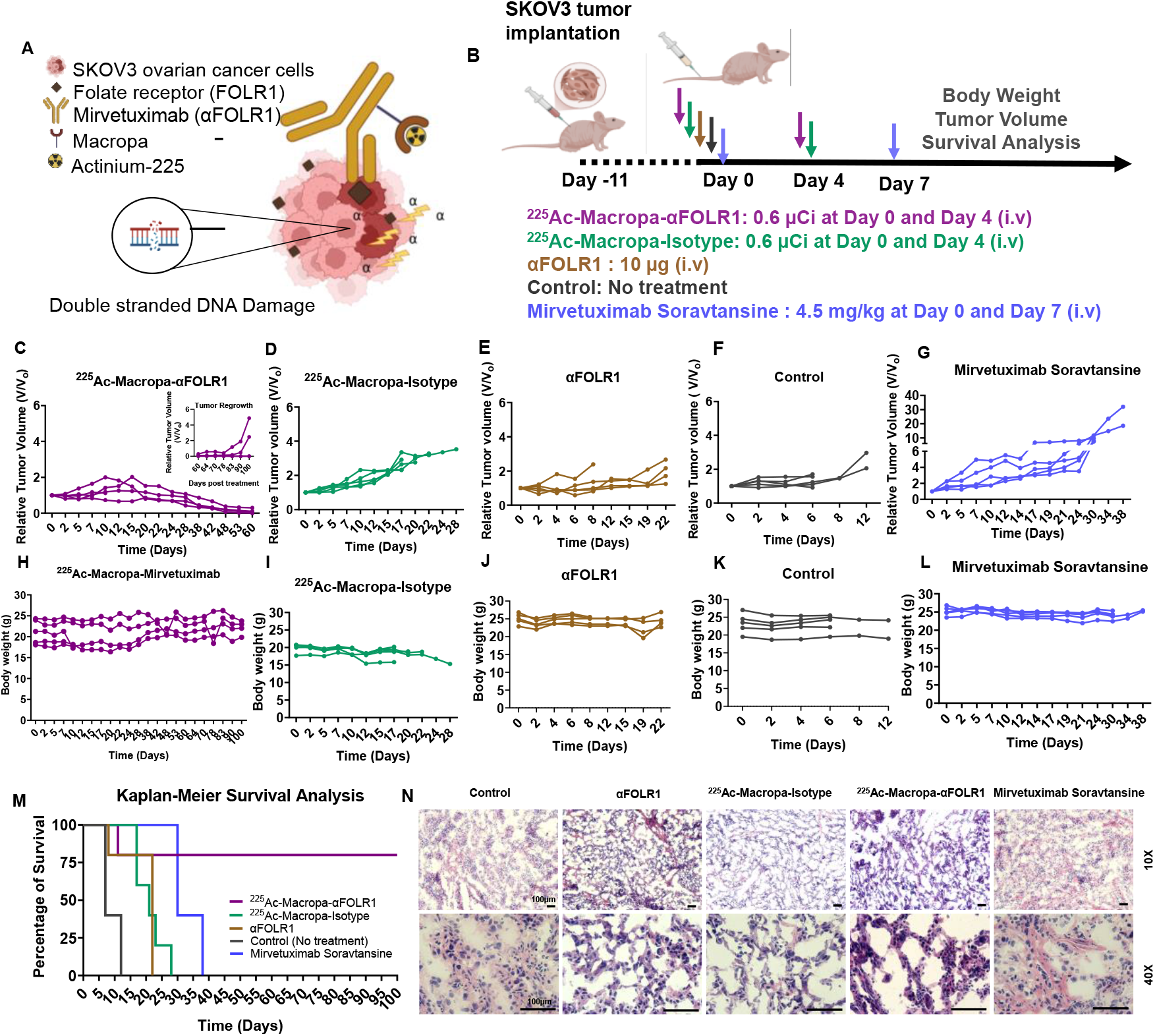
Evaluation of FOLR1-targeted Actinium-225 Therapy. (A) Schematic of mechanism of tumor killing by FOLR1 targeted actinium-225 therapy. (B) Dosing regimen for two-dose therapy. Relative tumor volume and body weight of (C, H) ^225^Ac-Macropa-αFOLR1, (D, I) ^225^Ac-Macropa-Isotype, (E, J) αFOLR1, (F, K) the blank control, and (G, L) Mirvetuximab Soravtansine (n=5). (M) Kaplan-Meier survival analysis. (N) Histological evaluation of tumor tissues at study end point by H&E (Magnification 10X (top panel) and 40X (bottom panel); Scale bar: 100µm).

Considering the rapid tumor retention and reduced blood uptake at 72 h p.i., we designed a two-step dosing regimen by administering a fractionated dose of 0.6 µCi per fraction (two doses, 4 days apart) (**Figure 5B**), to ensure sufficient tumor uptake and controlled blood kinetics. Treating with ^225^Ac-Macropa-αFOLR1, we observed a remarkable reduction in the relative tumor growth (**Figure 5C-F**) and tumor volumes (**Supplementary Figure 4 and 5**) compared to controls, including ^225^Ac-Macropa-Isotype (**Supplementary Figure 6)**, αFOLR1 alone and the blank control (n = 5 in each group). The fractionated dose was well tolerated as indicated by no significant changes in the body weight (**Figure 5H-K**). Only one mouse in the treated group showed a drop in body weight and eventual fatal response. Our findings indicated a 100% response rate; tumor regression in all ^225^Ac-Macropa-αFOLR1 treated mice was consistent for over 80 days (**Figure 5C**), which is more than 10% of the lifespan of mouse species. Complete tumor elimination was found in 40% of ^225^Ac-Macropa-αFOLR1 treated mice (**Supplementary Figure 5**) without any recurrence, while the other 40% of treated mice recurred after stopping treatment after 83 days. Since minimal radioresistance is expected in TAT, recurrence after 83 days (10% of the lifespan) will not be a major concern, and the exact same therapeutic protocol can be applied prior to recurrence to prevent tumor regrowth in the real clinical setting. Further, Kaplan-Meir survival analysis (**Figure 5M**) suggests an 80% survival rate for ^225^Ac-Macropa-αFOLR1 treated mice. The median survival for ^225^Ac-Macropa-αFOLR1 (100 days) was significantly higher than the controls (^225^Ac-Macropa-Isotype: 21 days; αFOLR1: 22 days; Blank control: 8 days). Such significant enhancement (10% of the lifespan of mice) of overall survival time is exciting, much longer than any of the current FDA-approved or in-trial therapeutics.

To better evaluate the clinical viability of our approach, we compared the treatment efficacy of ^225^Ac-Macropa-αFOLR1 against the FDA-approved antibody drug conjugate Mirvetuximab Soravtansine. Clinically, patients receive dose of Mirvetuximab Soravtansine once every 3 weeks until disease progression or unacceptable toxicity^16^. We administered two doses of Mirvetuximab Soravtansine at 4.5 mg/kg weekly^20^, the clinically recommended dose to match with our ^225^Ac-Macropa-αFOLR1 treatment. Although Mirvetuximab Soravtansine showed moderate therapeutic efficacy compared to blank control, it was not potent enough in SKOV-3 tumor bearing mice (**Figure 5G, Supplementary Figure 7**), with a median survival of only 30 days (**Figure 5M**), which is much shorter than what we achieved with ^225^Ac-Macropa-αFOLR1.We did not observe any obvious changes in mouse body weight (**Figure 5L**). Our results are consistent with the reported preclinical efficacy of Mirvetuximab Soravtansine in SKOV3 xenografts^20^. Despite targeting the same receptor and using the same ligand, TAT performed significantly better than targeted chemotherapy. Overall, our data establishes the superiority of ^225^Ac-Macropa-αFOLR1 which surpassed the therapeutic efficacy of Mirvetuximab Soravtansine in animal models. More clinically relevant models will be tested in the follow-up studies.

To confirm the therapeutic efficacy, hematoxylin and eosin (H&E) histological evaluation of tumor tissues was performed at the study end point (**Figure 5N**). Nuclei condensation was observed in ^225^Ac-Macropa-αFOLR1 treated mice in residual tumors, indicating strong cell apoptosis. Nuclear division was observed due to tumor regrowth after 83 days post treatment. Mice with complete tumor elimination had no residual tumor tissue for histological analysis.

*Ex vivo* biodistribution of the decay daughter Bismuth-213 (^213^Bi) was analyzed using gamma counter, to further validate the biodistribution of ^225^Ac-Macropa-αFOLR1 and control groups. We assessed the biodistribution of ^213^Bi-Macropa-αFOLR1 at 72h, after single dose of ^225^Ac-Macropa-αFOLR1 (0.6 µCi) (**Supplementary Figure 8)**. The tumor uptake of ^213^Bi-Macropa-αFOLR1 (16.95 ± 5.5 %ID/g) was significantly higher than the ^213^Bi-Macropa-Isotype (5.07 ± 0.41 %ID/g). The same study was repeated 14 days post treatment. The tumor uptake at Day 14 after two doses of ^225^Ac-Macropa-αFOLR1 (0.6 µCi, 4 days apart) was 127 ± 6.9 % ID/g while minimal activity observed in isotype control group. Of note, since the decay daughter ^213^Bi cannot be stably labeled, the quantification of ^213^Bi biodistribution may not perfectly match the ^89^Zr biodistribution in PET. However, the ^213^Bi biodistribution in our study further confirmed the strong tumor uptake in ^225^Ac-Macropa-αFOLR1 treated mice and demonstrated similar distribution pattern as PET imaging, which again attests to the superior tumor targeting of FOLR1 for effective TAT.

### ^225^Ac-Macropa-αFOLR1 causes tumor specific DNA double-strand breaks

One of the major advantages of alpha-particle therapy is its ability to inflict potent DNA double-strand breaks^9. 225^Ac decay generates four alpha particles with energies ranging from approximately 5.8 to 8.4 MeV, allowing a high linear energy transfer for DNA double-strand breaks^9,21,22^. Compared with conventional radiotherapy including X-ray and beta-particle therapies, alpha therapy can be more lethal to the tumor cells and less impacted by acquired radioresistance from DNA repair mechanisms^6,7^. To assess the tumor damage in this study, the tumor tissues from SKOV3 xenografts treated with 0.6 µCi (2 two doses, 4 days apart) ^225^Ac-Macropa-αFOLR1 were stained with H&E and γ-H2AX, a DNA damage marker for DNA double-strand breaks. The H&E images (**Figure 6A**) demonstrated deformed pyknotic nuclei in the ^225^Ac-Macropa-αFOLR1 treated group at both day 4 and day 14 post treatment compared to controls. Further, treatment with ^225^Ac-Macropa-αFOLR1 increased the γ-H2AX foci in the cell nucleus as early as Day 4 post treatment compared to ^225^Ac-Macropa-isotype, indicating early therapeutic effect. At day 14 post treatment, there was an increase in the DNA double strand breaks in ^225^Ac-Macropa-αFOLR1 treated group as indicated by the high intensity of the γ-H2AX signal and presence of karyorrhectic nuclei. This also coincides with an observed decrease in tumor volumes in the treated groups (**Figure 5C**). In all control groups, minimal γ-H2AX staining was observed confirming that ^225^Ac-Macropa-αFOLR1 treatment augments DNA damage in tumors.

**Figure 6.**
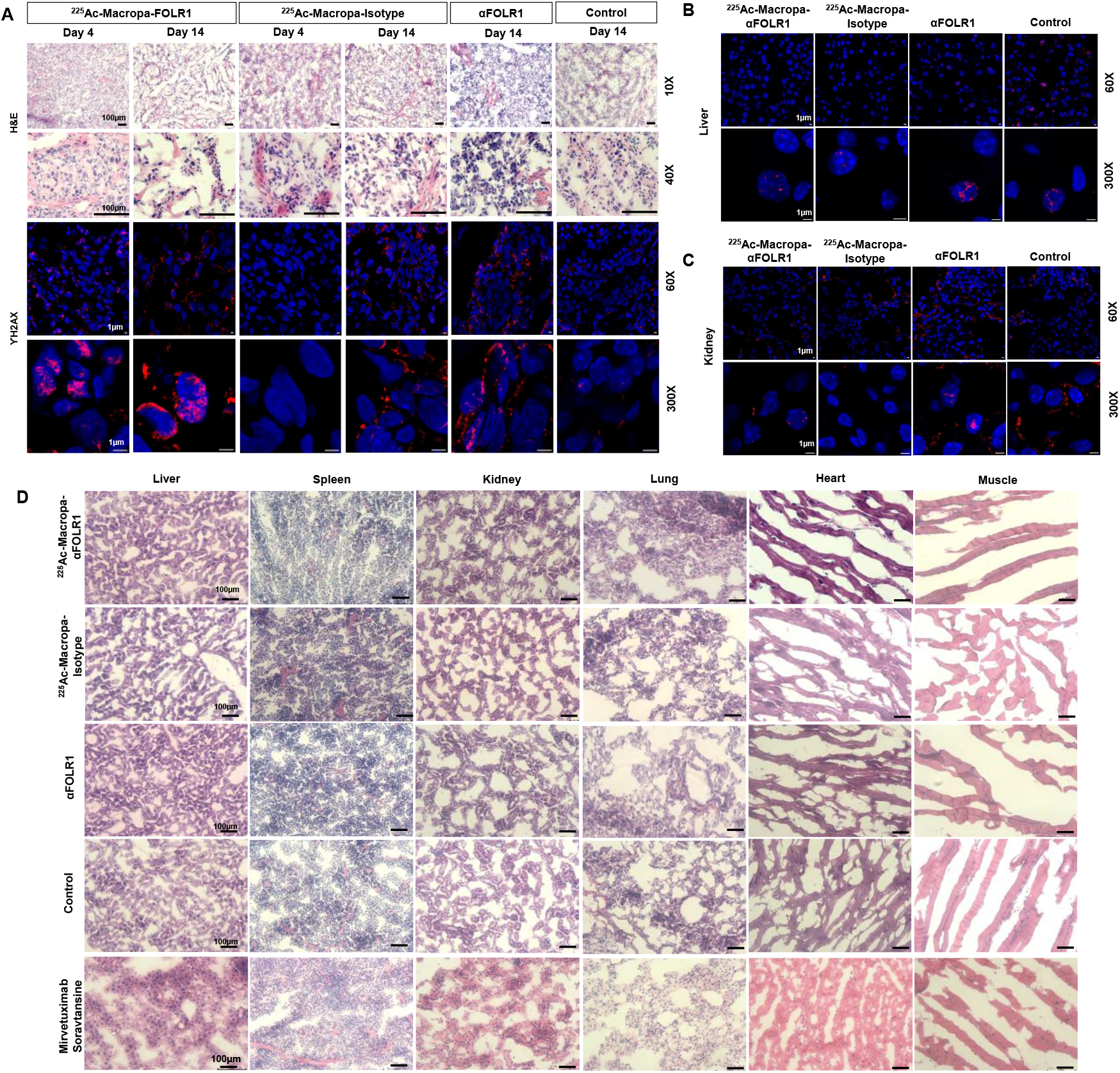
Evaluation of tumor killing and toxicity to normal healthy tissues. (A) Histological evaluation of tumor by H&E (top panel) and γ-H2AX double stranded DNA damage marker (bottom panel) at Day 4 and Day 14 for ^225^Ac-Macropa-αFOLR1 and ^225^Ac-Macropa-Isotype with control and αFOLR1 evaluated at Day 14 (Images acquired at magnification 10X and 40X for H&E; Scalebar: 100µm). γ-H2AX staining for (B) liver and (C) Kidney tissues at Day 14 post treatment (For γ-H2AX, images acquired at magnification 60X and 60X Zoom 5, Scale bar: 1µm). (D) Histological evaluation of normal tissues at study end point by H&E staining for ^225^Ac-Macropa-αFOLR1, ^225^Ac-Macropa-Isotype, αFOLR1, blank control and Mirvetuximab Soravtansine.

### Histological evaluation exhibits no obvious toxicity in normal tissues

Cytotoxicity is one of the major concerns of TAT, considering the high linear energy transfer of the radioisotopes which can create lethal DNA damage in any recipient cell, no matter whether it is tumor or normal tissue. Although we have successfully demonstrated that FOLR1-based targeting has significantly enhanced tumor uptake with minimal nonspecific accumulation in normal tissues, it is still critical to investigate the histological toxicity in off-target organs. Thus, we assessed DNA double-strand breaks in the clearance organs, liver and kidney, by γ-H2AX staining at Day 14 post treatment. As shown in (**Figure 6 B and C**), we did not observe any prominent γ-H2AX foci in the cell nucleus of liver and kidney, demonstrating the tumor-specificity and safety of the treatment. Besides, H&E evaluation indicated no obvious changes in the structural integrity of tissues at Day 14 (**Supplementary Figure 9**) and the study end point (**Figure 6D**). Overall, we confirmed that by sophisticated design of the treatment plan and dosimetry (a two-step treatment with 0.6 µCi per dose in this study), TAT can be very safe without inducing any major radiotoxicity.

### Radiation dosimetry reveals dominating tumor accumulation

Since this is the first study of ^225^Ac-based FOLR1-targeted alpha therapy in ovarian cancer and there was no preliminary reference for dosimetry of related agents, we performed intensive dosimetry analysis. These assessments are critical to provide important guidance for future studies, including radiation dose quantification, prediction of therapeutic effects and toxicity tolerance, and minimizing animal use. The dosimetry data in this study can also be useful for guiding future clinical translation in humans. Radiation dosimetry calculations were performed by fitting the biodistribution data for tumor and normal tissues using MIRDfit^23^ and estimating the whole-body absorbed doses per injected activity in the MIRDcal software^24,25^. The biodistribution data for all non-malignant organs fitted well using monoexponential biokinetic function with R^2^>0.95 except for the liver (R^2^ = 0.72) in the ^225^Ac-Macropa-Isotype group (**Supplementary Figure 10**,**11**). Biexponential fitting of tumor biodistribution data resulted in R^2^ of 0.95 (**Supplementary Figure 11G**,**12G**). The ^225^Ac-Macropa-αFOLR1 exhibited high time integrated activity curve (TIAC) in tumors (60.7 h) compared to healthy organs and tissues (**Supplementary Table 1**). The TIAC for ^225^Ac-Macropa-Isotype was 2.8 h in tumors whereas liver showed a high TIAC of 45 h (**Supplementary Table 2**).

The tumor tissue showed the highest absorbed dose of 2.43 ± 0.362 Gy/MBq in ^225^Ac-Macropa-αFOLR1 group (**Figure 7A, Supplementary Table 3**), compared to 0.139 ± 0.144 Gy/MBq by ^225^Ac-Macropa-Isotype. The calculated total absorbed dose for ^225^Ac-Macropa-αFOLR1 (kidney: 9.4 ± 2.7 mGy/MBq, spleen: 22.9 ± 8.7 mGy/MBq, liver: 24.9 ± 7.3 mGy/MBq) was minimal in all healthy organs compared to ^225^Ac-Macropa-Isotype (kidney: 16.6 ±7.9 mGy/MBq, spleen: 59 ± 47.7 mGy/MBq, liver: 87.2 ± 72.2 mGy/MBq). In addition, ^225^Ac-Macropa-αFOLR1 showed a high tumor-to-muscle (1278 ± 792), tumor-to-heart (1664.3 ± 940.2), tumor-to-liver (97.6 ± 50) and tumor-to-kidney (258.5 ± 129.7) ratio in comparison to ^225^Ac-Macropa-Isotype (**Figure 7B**).

**Figure 7.**
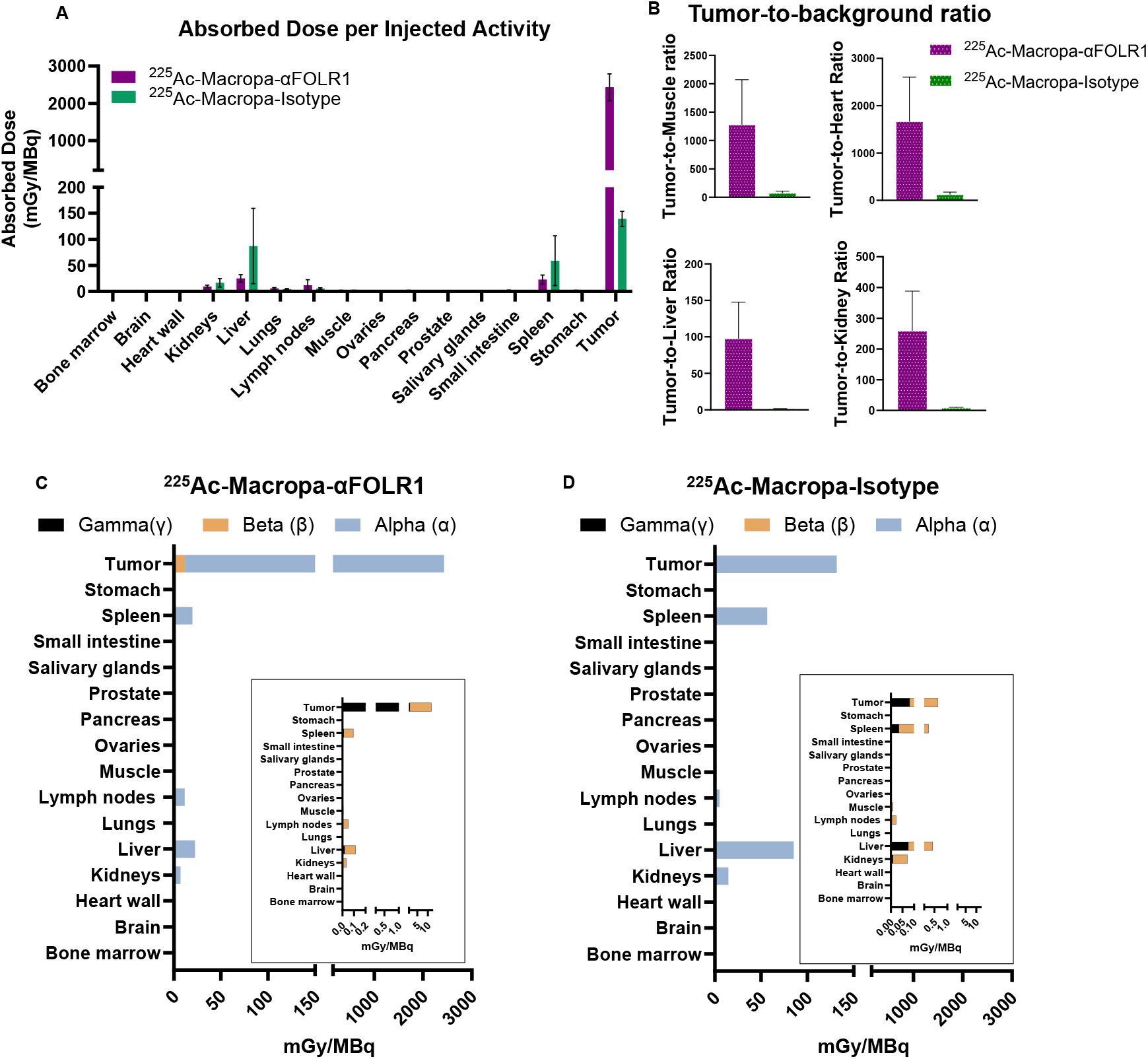
Estimated human dosimetry of ^225^Ac-Macropa-αFOLR1. (A) Total absorbed dose calculations per injected activity for ^225^Ac-Macropa-FOLR1 and ^225^Ac-Macropa-Isotype (B) Absorbed dose ratio for tumor-to-muscle, tumor-to-heart, tumor-to-liver, and tumor-to-kidney. Contribution of alpha (α), beta (β), and gamma (γ) to absorbed self-dose of (C) ^225^Ac-Macropa-FOLR1 and (D) ^225^Ac-Macropa-Isotype.

Further, most of the total absorbed was from alpha particles (**Figure 7C, D; Supplementary Table 4**,**5**). The tumors demonstrated pronounced alpha-particle self-absorbed dose from ^225^Ac-Macropa-αFOLR1. Gamma ray and beta-particle absorbed dose (**Figure 7C, D**) was minimal compared to the alpha-particle dose. In the control group, the gamma ray and beta-particle absorbed dose contribution was higher than ^225^Ac-Macropa-αFOLR1 but remained considerably low compared to alpha-particle self-absorbed dose. Further, the cross absorbed dose (**Supplementary Figure 12**) from alpha particles in ^225^Ac-Macropa-αFOLR1 was under 6% in all organs comparable to ^225^Ac-Macropa-Isotype.

## Discussion

Surgery and platinum-based chemotherapy remain the cornerstone of treatment of ovarian cancers. However, inherent and acquired drug resistance is inevitable and poses a major challenge in sustainable treatment of these cancers. Recent FDA approval of Mirvetuximab Soravtansine has shown improved therapeutic responses in patients with platinum-resistant ovarian cancer. Though the antibody-drug conjugate demonstrates superior specificity and anti-tumor effects, acquired resistance with repeated use has been observed in patients^26,27^. Moreover, although the antibody-drug conjugate increases survival by 4 months, there remains much room for improvement. Targeted radionuclide therapies, in particular alpha-particle therapy, has demonstrated excellent anti-tumor efficacy compared to chemotherapy and conventional radiotherapy including beta particle-based treatments^28,29^. The high linear energy transfer (50 to 230 keV/µm) enables alpha particles to induce irreparable double stranded DNA breaks at the target site. These cytotoxic effects by alpha particles are independent of oxygen free radicals and do not depend on the phase of the cell cycle^30^. These properties make TAT an attractive strategy for recalcitrant cancers such as ovarian and other gynecological malignancies.

The U.S. Department of Energy (DoE) highlighted the potential applications of ^225^Ac in 2018, opening a new era of alpha particle-based molecularly targeted therapy for cancer. The National Cancer Institute (NCI) has designated staunch support for R&D and production of alpha emitters for cancer therapy. Among all the alpha emitters, ^225^Ac is the most promising because of its suitable half-life (∼9.9 days) and high emission density (the emission of four alpha particles in its decay chain). Therefore, developing targeted radiotherapy with ^225^Ac and exploring its applications in hard-to-treat cancers is timely and highly innovative.

Considering the highly localized energy deposition and strong biological impact of alpha particles, tumor-selective delivery is of paramount importance. Although previous studies have evaluated CD276^31^, tumor-associated glycoprotein 72 (TAG-72)^32^, HER2^33-36^, HER3^37^, and MUC16^38^ as targets for antibody-based alpha therapy development in ovarian cancer, FOLR1 has gathered considerable interest as a therapeutic target in ovarian cancer. Our transcriptome analysis of clinical datasets reinforced the significantly high expression of FOLR1 in ovarian cancers compared to healthy ovary tissue and other cancer types. Our findings indicated no notable change in the FOLR1 expression at distinct stages of ovarian cancer. Therefore, FOLR1 targeting with ^225^Ac-Macropa-αFOLR1 can be a widely applicable agent for TAT in majority of patients with ovarian cancer, irrespective of disease stage or tumor burden.

As determined by transcriptome data and immunofluorescence, the SKOV3 model demonstrated moderate FOLR1 expression and thus can be representative of response in both high and medium FOLR1 expressing tumors. Indeed, our data showed that ^89^Zr-DFO-αFOLR1 exhibited tumor-specific uptake with minimal accumulation in other organs when compared to ^89^Zr-DFO-Isotype control. This tumor uptake was remarkably high and exceeded that of other published work for ovarian cancers with standard radiolabeled antibodies targeting HER2, MUC16, TAG-72^32-36^ following intravenous administration, further underscoring the specificity of αFOLR1 to deliver ^225^Ac at the tumor site.

Our studies further demonstrated that Macropa-based ^225^Ac labeling of the antibodies was ultra stable, and despite differences in chelator-radioisotope constructs, PET imaging can truly represent the biodistribution of our tracers. We have proven that both ^89^Zr-DFO-αFOLR1 and ^225^Ac-Macropa-labeled antibody can be stable up to at least 72 h in both PBS buffer and mouse serum. This period is sufficient for antibody tracer to clear from the bloodstream and primarily accumulate in the tumors, reducing concerns of potential detachment or toxicity of ^225^Ac. Therefore, we are confident that the PET imaging can truly represent the real biodistribution of ^225^Ac-labeled antibody, and changing the isotopes and chelators will not significantly impact its pharmacokinetics. Moreover, Macropa has shown superb advantages over traditional chelators, including room temperature chelation, excellent stability, and high-fidelity chelation for decay daughters. Recent findings indicate the ability of Macropa to recapture the decay daughter ^213^Bi in situ addressing the long-standing issues of non-specific radiotoxic effects of free ^213^Bi^39^. Our own results in this study corroborate these findings. Overall, our work sets an important foundation for future studies, using ^89^Zr-DFO-based PET imaging to evaluate biodistribution and predict the dosimetry for ^225^Ac-Macropa-based TAT.

We report the first FOLR1 targeted ^225^Ac therapy for treatment of ovarian cancers with exceptional tumor growth control and extended survival rates. Two fractionated doses of 0.6 µCi (22 kBq) of ^225^Ac-Macropa-αFOLR1 were well tolerated, achieved a high ∼80% survival rate and prolonged animal survival for 60-90 days compared to controls (12-28 days). *Ex vivo* biodistribution at day 14 after fractionated two dose therapy indicated significantly higher tumor uptake of ^225^Ac-Macropa-αFOLR1 in tumors compared to ^225^Ac-Macropa-Isotype, which is the highest dose (44 kBq ^225^Ac) reported so far without causing obvious toxic effects. Our study reported a 40% complete tumor regression, which is higher than FDA-approved and in-trial therapies. The other 40% of mice showed recurrence after stopping treatment for 83 days. These recurrences can be possibly attributed to residual cancer cells that transition from dormant state to proliferative state^40,41^. At molecular level, tumor relapses are regulated by numerous factors such as genetic and epigenetic changes, signaling pathways, tumor microenvironmental factors such metabolic adaptations, immune system changes and presence of cancer stem cells^42^. Considering the fact that FOLR1-based targeting is very specific with tolerable toxicity and TAT is less susceptible to traditional radioresistance mechanisms, more treatment cycles can be applied to prevent tumor recurrence without increasing side effects. Exploring combination strategies with immune checkpoint inhibitors, PARP inhibitors or β emitters can further potentiate the therapy outcomes of TAT^43^. Of note, this proof-of-concept study is the first study of FOLR1-based TAT in ovarian cancer and only single tumor model SKOV3 was evaluated, which may be far from the real clinical scenario. Future studies will replicate the treatment in more tumor models, including tumors in immune competent mice and genetically engineered mice, in our follow-up studies to validate and optimize the concept, methodology and protocols developed in this study.

This study provides one of the first dosimetry references for FOLR1-based TAT. Using the Medical Internal Radiation Dose (MIRD) standard dosimetry method, we proved that the tumor tissue received the highest absorbed dose of 2.43 ± 0.362 Gy/MBq in ^225^Ac-Macropa-αFOLR1 group, compared to 0.139 ± 0.144 Gy/MBq by ^225^Ac-Macropa-Isotype. This extreme high exposure confirms the targeting specificity of FOLR1-targeted antibody, which is effective to eliminate ovarian cancer in animal models. Follow-up studies will be performed by varying treatment doses, to explore the minimal effective dose that can cure tumors without obvious toxicity.

In summary, we present a novel ^225^Ac-based FOLR1-targeted alpha-particle therapy for treatment of ovarian cancer. Proven by noninvasive and quantitative PET imaging in a preclinical animal model, αFOLR1 exhibited pronounced tumor specific uptake and reasonable biodistribution profile, ideal for delivering high energy ^225^Ac for TAT. FOLR1-based TAT exhibited enhanced antitumor efficacy, survival, tolerability, and favorable radiation dosimetry. Considering the extraordinary therapeutic efficacy and the fact that all the components, including the αFOLR1 antibody and the isotope ^225^Ac, are clinically approved or in-trial, our approach has promising potential for clinical translation. We envision that FOLR1-based TAT will eventually benefit patients suffering from ovarian cancer to battle against this common but lethal disease.

## Methods

### Analysis of FOLR1 expression

Transcriptomic data from the Cancer Cell Line Encyclopedia (CCLE) was accessed from a publicly available website. Gene expression levels (log_2_ [TPM + 1]) were extracted and compared across tumor lineages. All plots were generated using R (v4.3.1) and ggplot2.

Transcriptomic data for tumor tissues and matched clinical annotations were obtained from The Cancer Genome Atlas (TCGA). Normal tissue data were retrieved from the GTEx project to supplement tumor-normal tissue comparisons in cancer types lacking paired adjacent tissues. Only TPM-normalized RNA-seq data were used in all analyses. Pan-cancer comparisons of FOLR1 expression were performed across TCGA tumor types, while tumor vs. normal analyses relied on non-paired comparisons with GTEx tissues. For ovarian cancer (OV), associations between FOLR1 expression and clinical stage, tumor characteristics, and survival outcomes (overall survival (OS), Disease specific survival (DSS), PFI) were examined. Statistical comparisons were conducted using the Wilcoxon rank-sum test for group differences and Kaplan– Meier analyses for survival.

### Cell culture and *vitro* assays

SKOV3 (wildtype) cells were maintained at 37 °C under 5% CO_2_ in McCoy’s 5A medium supplemented with 10% fetal bovine serum and 1% penicillin-streptomycin. Before the initiation of experiments, cells were maintained at ≤ 75% confluency for at least 72 h by routine sub-culturing involving trypsinization, centrifugation, and re-seeding in T75 culture flasks.

#### Assessment of FOLR1 expression

The FOLR1 expression was assessed using FOLR1 targeted antibody (Mirvetuximab) by flow cytometry. Mirvetuximab (αFOLR1) (MedChem Express) and Human IgG1 (isotype control) (BioXcell) were conjugated with Alexa fluor 488 at 1:10 molar ratio for 2 h at room temperature and pH 8.5. The free Alexa fluor 488 was purified by PD10 columns (Cytiva) using phosphate buffer saline as the elution buffer. SKOV3 cells were cultured in folate free media for at least two passages and were harvested and suspended in phosphate-buffered saline (PBS) supplemented with 2% fetal bovine serum (FBS) at a concentration of 5 × 10^6^ cells/mL. Cells were incubated with Alexafluor 488-labeled αFOLR1 and isotype conjugates (5µg equivalent to Alexafluor 488 concentration) for 1 h at 24°C. After washing twice, samples were analyzed with a Cytoflex LX (University of Utah Flow core). FlowJo software (v10.10) was employed for analysis of flow cytometry data (Tree Star Inc.)

### Animal models

Female athymic nu/nu mice (4-6 weeks old) from Charles River Laboratories were housed under sterile conditions. Animal care and procedure were conducted with Institutional Animal Care and Use Committee (IACUC) approval, incompliance with all federal and institutional guidelines for the care and use of the laboratory animals at the University of Utah. Female athymic nu/nu mice were injected subcutaneously with 3×10^6^ SKOV3 (wildtype) ovarian cancer cells mixed with Matrigel at 1:1 ratio. The body weight and tumor volume [0.5(length × width^2^)] were measured.

### Radiolabeling and Stability

#### Radiolabeling with ^89^Zr

αFOLR1 and Human IgG1 were reacted with p-SCN-Bn-Deferoxamine (DFO) (Macrocyclics) in a 1:10 molar ratio at pH 8.5-9 at room temperature for 2 h. The reaction was purified by PD10 columns using 0.5M HEPES (pH 7) as the elution buffer. The DFO conjugated αFOLR1 and Human IgG1 (100-200 ug) were mixed with ^89^Zr oxalate solution (Washington University, St. Louis) in HEPES buffer (0.5M, pH 7) and was performed at 37°C for 1 h. The progression of labeling reaction was monitored with iTLC (Agilent Technologies, Inc) at 0, 15, 30, 45 and 60 minutes using EDTA (50 mM, pH 7) as the mobile phase. The free ^89^Zr was purified using PD10 columns. The labeling efficiency and radiochemical purity were assessed by iTLC, and radioactivity was measured on gamma counter.

#### Radiolabeling with ^225^Ac

αFOLR1 and Human IgG1 were reacted with Macropa-Bz-NCS (Ratio Therapeutics) at a 1:5 molar ratio for 2h (pH 8.5-9) at room temperature. The free chelator Macropa-Bz-NCS was purified using PD10 columns using ammonium acetate (0.1 M, pH 7) as the elution buffer. For generation of Actinium-225 antibody conjugates, ^225^Ac nitrate solution (15-20 µCi) obtained from the National Isotope Development Center (NIDC) was added to ammonium acetate (0.1 M, pH 5.5) buffer and mixed with Macropa conjugated αFOLR1 and Human IgG1 (20-40 ug). The pH was adjusted to 5.5 and reaction was carried out for 2 h at room temperature. The labeling efficiency and radiochemical purity were assessed by iTLC. The radioactivity on iTLC was measured on a gamma counter. The percentage of radionuclide was calculated as

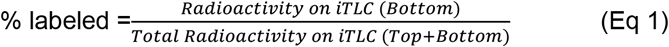

#### Stability studies

*In vitro* stability of ^89^Zr-DFO-αFOLR1, ^89^Zr-DFO-Isotype, ^225^Ac-Macropa-αFOLR1 and ^225^Ac-Macropa-Isotype control was assessed in serum and phosphate buffer saline for 72 h. For the stability studies, 30 μCi ^89^Zr-DFO-mAb or 0.1 µCi ^225^Ac-Macropa-mAb was added to 100 μL of 100% mouse serum or phosphate buffer saline and the mixture was incubated at room temperature. Aliquots were collected at different points: 0, 6, 24, 48 and 72 h. The stability was assessed by iTLC with EDTA as the mobile phase. The iTLC plates were cut in half and radioactivity was measured from top-to bottom-half portions of iTLC using gamma counter. The percentage of radionuclide labeled to the antibody was calculated using equation 1.

### *In vivo* PET imaging, Pharmacokinetics and Biodistribution

Five to seven weeks old athymic nu/nu mice bearing subcutaneous SKOV3 tumors were injected intravenously with radiolabeled mAbs, and PET images were acquired at 24, 48, 72 h on Siemens Inveon PET scanner. After the final scans at 72 h, the mice were euthanized to collect blood and tissues. Tissues were weighed, and radioactivity was measured on a gamma counter (Perkin Elmer). Final values were decay corrected, and results are expressed as a percentage of injected dose per gram of tissue (%ID/g). For image quantitation, PET images were reconstructed, and ROIs were drawn over organs of interests on Inveon acquisition workplace 2.1. For pharmacokinetic analysis, PET signal (%ID/g) was plotted over time to generate time-activity curves for the main organs of interest.

For the biodistribution of ^225^Ac-Macropa-αFOLR1 and ^225^Ac-Macropa-Isotype, a single dose of 1 µCi or two doses (0.6 µCi each, 4 days apart) was injected intravenously, and the mice were euthanized on day 14. The organs were harvested, weighed and remaining ^225^Ac radioactivity was measured by assessing the activity of the ^213^Bis window in a gamma counter. In another study, mice were injected with 0.6 µCi of ^225^Ac-Macropa-mAbs and the *ex vivo* biodistribution of ^225^Ac radioactivity of ^213^Bis was assessed at 72 h by a gamma counter.

### *In vivo* therapeutic efficacy in SKOV3 Xenografts

The antitumor efficacy of ^225^Ac-Macropa-αFOLR1 was evaluated as a monotherapy in the FOLR1 expressing SKOV3 subcutaneous tumor. When the tumors reached 50-100 mm^3^ volume, a single dose of 1 µCi of ^225^Ac-Macropa-αFOLR1 and ^225^Ac-Macropa-Isotype was intravenously administered. For evaluation of *in vivo* antitumor efficacy of two dose therapy, the mice were grouped into ^225^Ac-Macropa-αFOLR1 (0.6 µCi at Day 0 and Day 4), ^225^Ac-Macropa-Isotype (0.6 µCi at Day 0 and Day 4), αFOLR1 (10 µg at Day 0) and no treatment control. Subcutaneous tumor growth was monitored by measuring tumor volume (0.5 × length × width^2^) using a vernier caliper. Animal body weight was monitored as an indicator of treatment-related toxicity. Measurement of tumor volume and body weight was performed two to three times per week. Individual animals were sacrificed when showing >20% body weight loss or when tumors reached 400 % of the original tumor volume or as recommended by animal care staff.

### Evaluation of toxicity

Tissues were harvested at the end point for therapy and biodistribution studies and embedded in an OCT tissue freezing medium. Tissues were cryo-sectioned into 5 µm thick sections (ARUP Laboratories, University of Utah) and stored at -80°C for further use. For histological evaluation of tissues for signs of toxicity, the sections were stained with hematoxylin and eosin (H&E), and images were acquired using a brightfield microscope (Amscope). For immunofluorescence, tissue sections were fixed with cold 4% PFA for 10 min and washed twice in PBS and blocked using blocking buffer (5% BSA in PBS supplemented with 0.1 % Tween 20) for 1 hour. For γ-H2AX staining, tissues were first incubated overnight at 4°C with primary rabbit γ-H2AX (1:100, diluted in blocking buffer). After washing twice with PBS, the sections were incubated for 1 hour at room temperature with secondary Donkey anti-Rabbit IgG Dylight 594 (1:200, diluted in blocking buffer). Sections were washed twice in PBS, stained with 300 nM DAPI for 10 minutes, followed by antifade mounting media. All slides were stored at 4°C in a light-protected environment. Fluorescence images were acquired with the confocal laser scanning microscope (CLSM, Leica SP8).

### Radiation dosimetry

Dosimetry analysis was performed using the MIRDCal software^24^. The biodistribution data were converted to percent injected dose (%ID) by multiplying the injected dose per g (%ID/g) by the weights of organs from the athymic nude mice cohort. We estimated normal-tissue biokinetics based on the average percent injected dose (%ID) values from the PET imaging data of ^89^Zr-DFO-αFOLR1 at 24, 48 and 72h.Time integrated activity curves (TIAC) were generated and fitted using the monoexponential (biologic clearance with physical decay) function in the MIRDfit software^23^.

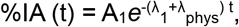

where A_1_ is the exponential amplitude representing the initial activity at time t = 0; λ_1_ is biological clearance constant (h^-1^); and λ_phys_ is the physical decay constant (h^−1^).

The estimated human organ-absorbed radiation doses per injected activity of ^225^Ac were calculated in MIRDcalc using the ICRP Adult Female phantomand the TIAC values obtained from MIRDcalc^25^.

For tumor dosimetry, we used hybrid approach integrating the average percent injected dose (%ID) values from the PET imaging data of ^89^Zr-DFO-αFOLR1 obtained at 24, 48, 72 h and biodistribution data of ^225^Ac-Macropa-αFOLR1 at 336 h. The time integrated activity curves for tumors were generated using biexponential fitting function.

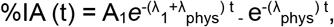

Where *A*_1_, exponential amplitude coefficients (%); λ_1_, λ_2_ = biologic clearance constants (h^−1^); λ_phys_ = physical decay constant (h^−1^).

Absorbed radiation dose calculations for tumors were performed using the integrated sphere tumor model in MIRDcalc.

### Statistical analysis

Data was analyzed using unpaired t-tests and two-way ANOVA (multiple comparisons) using GraphPad Prism 10. Statistical significance between different experimental groups was measured at unpaired t test and two-way ANOVA p-values of * p < 0.05, ** p < 0.01, ** p < 0.001 and *** p < 0.0001.

## Acknowledgements

This work was supported, in part, by the University of Utah School of Medicine (S.S), University of Utah College of Pharmacy (S.G.), the University of Utah Immunology, Inflammation and Infectious Diseases (3i) Initiative (S.G. and S.S.), the 5 For the Fight Fellowship (S.G.), and the Office of the Vice President for Research Seed Grants (S.S.). We acknowledge direct financial support from the Huntsman Cancer Center supported by the National Cancer Institute of the NIH under award number P30CA042014. We acknowledge Ratio Therapeutics for providing Macropa. The content is solely the responsibility of the authors and does not necessarily represent the official views of the NIH.

## Author contributions

N.S. performed all the experiments and data analysis and wrote the manuscript. E.N., A.B. and M.G. assisted in animal studies. L.W. and F.M. performed dosimetry analysis and assisted in manuscript writing related to dosimetry. F.G. performed transcriptome analysis. T.M. provided guidance on alpha-particle radiochemistry and edited the related part in the manuscript. T.A and A.K.S. provided guidance on ovarian cancer biology and edited the related part in the manuscript. A.M provided guidance on radiotherapy and edited the related part in the manuscript. S.C.M. provided guidance on radiobiology and toxicity and edited the related part in the manuscript. S.M. provided guidance on molecular imaging and edited the related part in the manuscript. S.G. and S.S. conceived and designed the experiments, oversaw and guided the whole study, and edited the manuscript.

