## Supplementary Information for "FOLR1-targeted Actinium-225-based Alpha-particle Therapy Eliminates Ovarian Cancer"

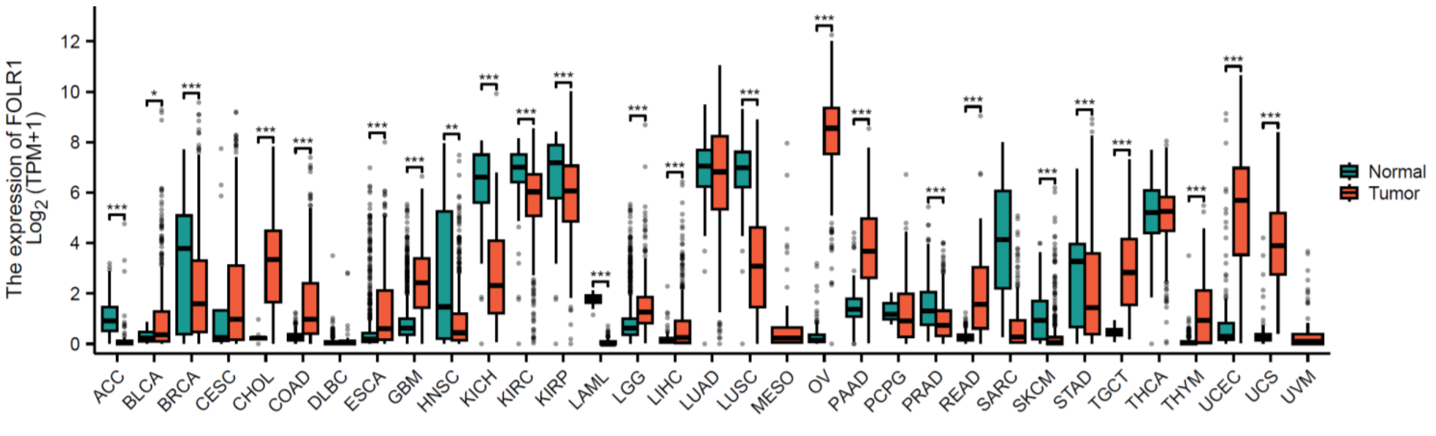


**S1. Expression of FOLR1 in cancer and normal tissues.** Gene expression levels of FOLR1 ($\log_2$ [TPM + 1]) across tumor lineage indicating highest expression in ovarian/fallopian cancers with minimal expression healthy ovary tissues.

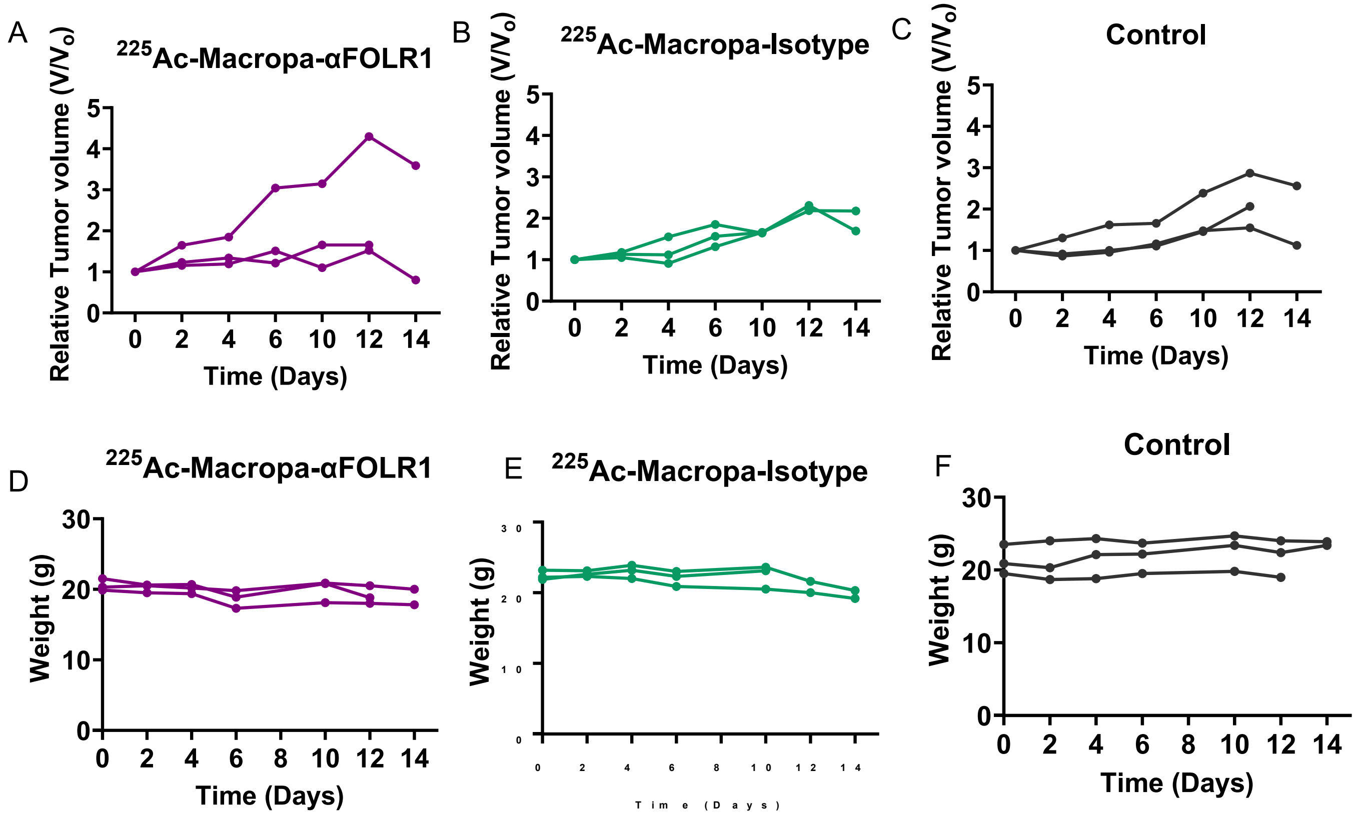


**S2. Therapeutic efficacy of single high dose therapy of $^{225}$Ac-Macropa-αFOLR1.** Relative tumor volume in SKOV3 subcutaneous tumor model treated with (A) $^{225}$Ac-Macropa-αFOLR1 (1 μCi) (B) $^{225}$Ac-Macropa-Isotype (1 μCi) and (C) Control (No treatment) group. Body weight for (D) $^{225}$Ac-Macropa-αFOLR1 (1 μCi) (E) $^{225}$Ac-Macropa-Isotype (1 μCi) and (F) Control (No treatment) groups. (n=3)

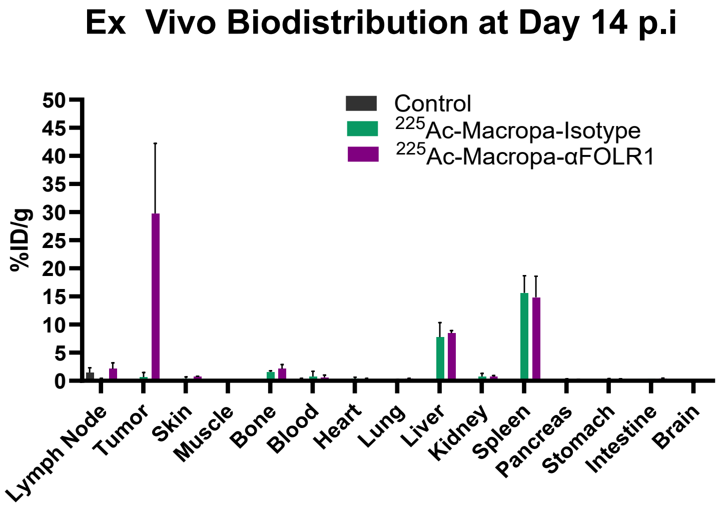


**S3. Ex vivo biodistribution of single high dose therapy of $^{225}$Ac-Macropa-αFOLR1** Radioactivity assessed by gamma counting at D14 in SKOV3 administered with single high dose of $^{225}$Ac-Macropa-αFOLR1(1 μCi) compared to $^{225}$Ac-Macropa-αFOLR1(1 μCi)

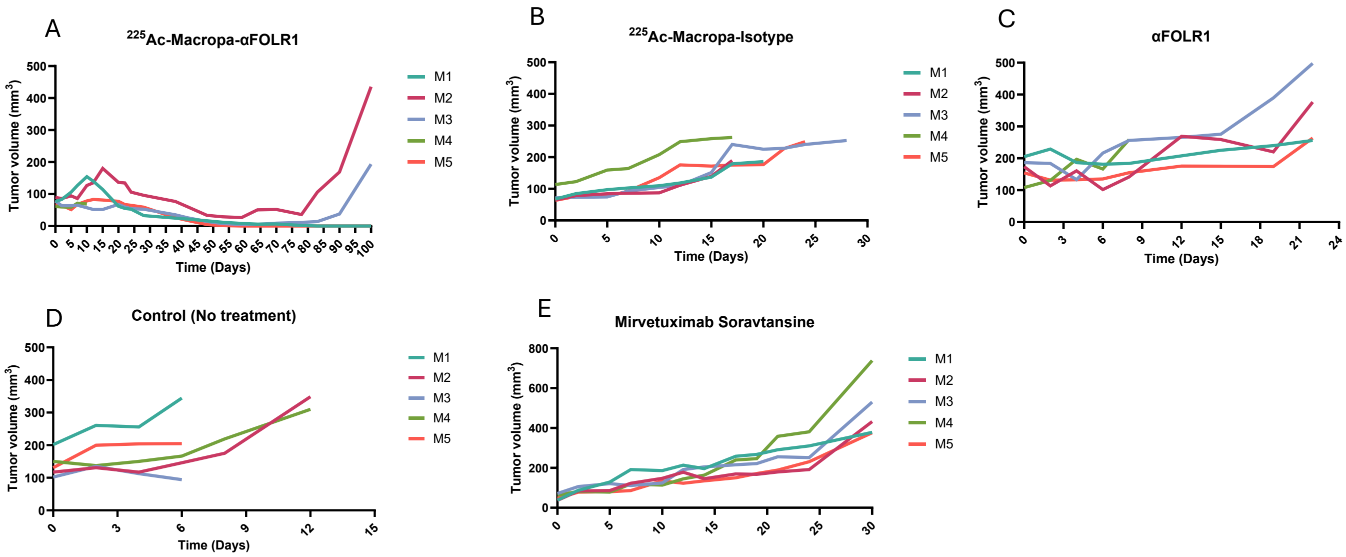


**S4. Tumor response with fractionated two dose therapy**. Tumor growth curves of SKOV3 xenografts treated with (A) $^{225}$Ac-Macropa-αFOLR1 (B) $^{225}$Ac-Macropa-Isotype (C) αFOLR1 (D) Control (no treatment) (E) Mirvetuximab Soravtansine (n=5). Some of the mice in the control groups were euthanized before reaching the tumor size limit, because they developed severe ulceration.

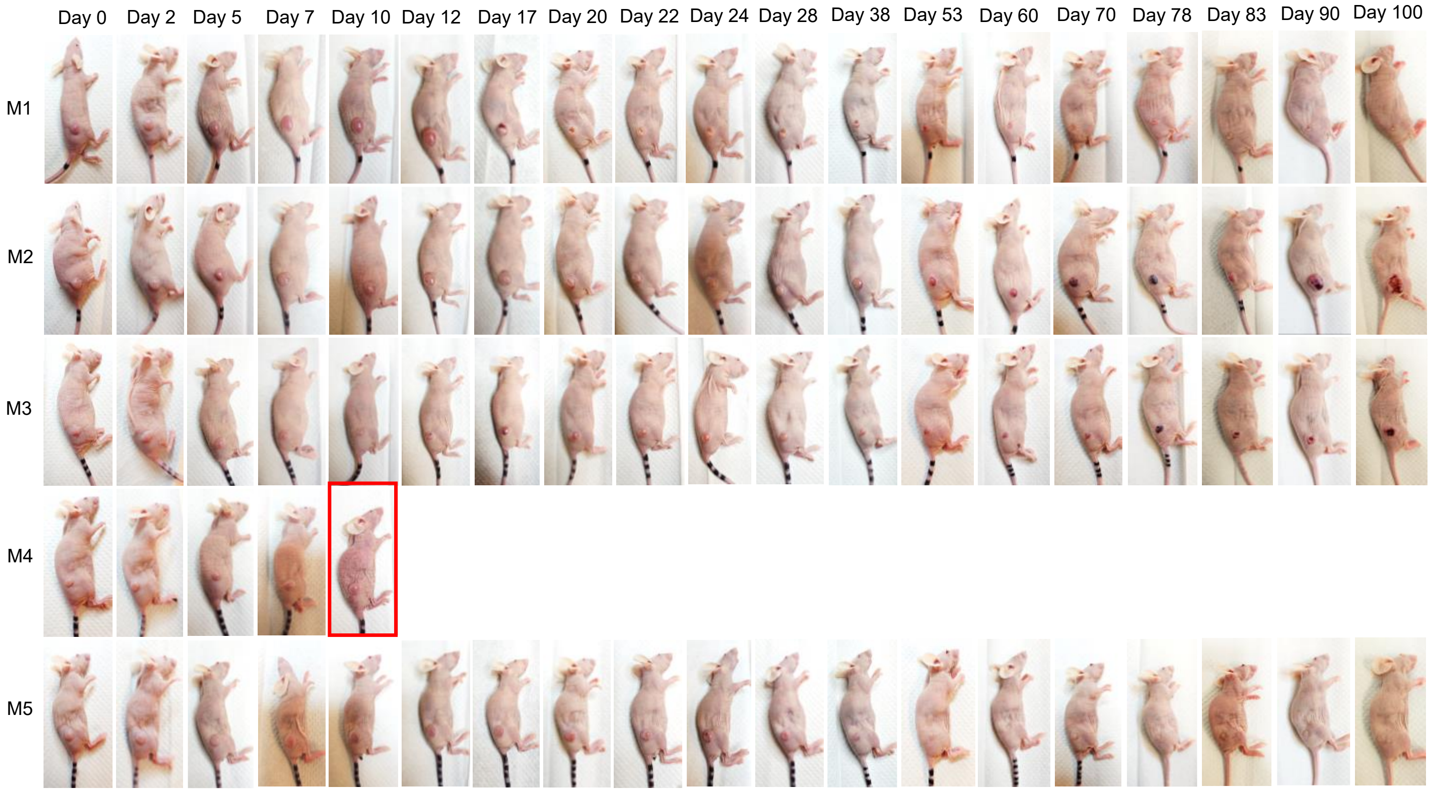


**S5. Tumor monitoring of $^{225}$Ac-Macropa- αFOLR1**. Visual assessment of tumor growth in SKOV3 xenografts administered with two doses (0.6 µCi, 4 days apart) of $^{225}$Ac-Macropa- αFOLR1.

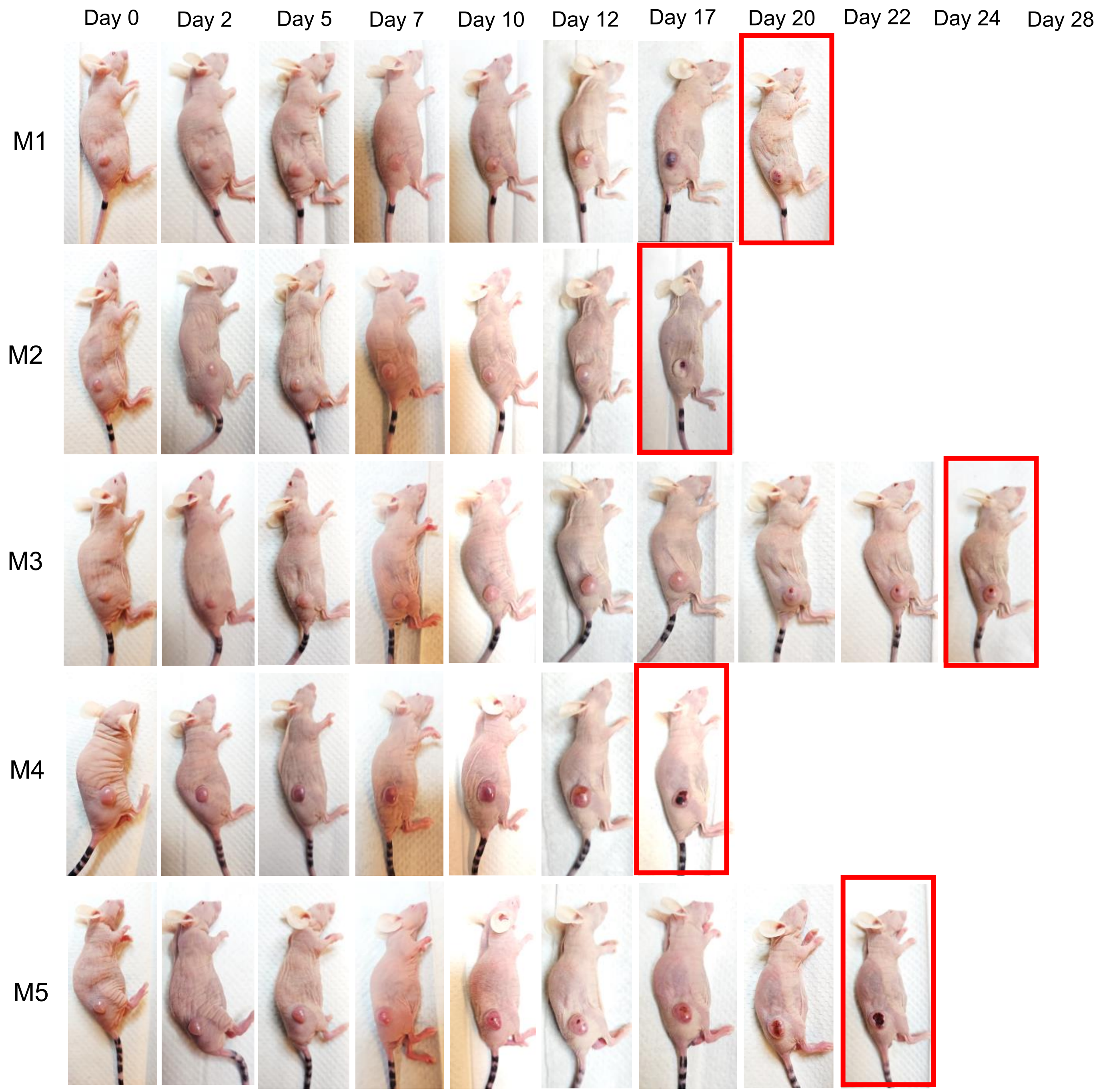


**S6. Tumor monitoring of $^{225}$Ac-Macropa-Isotype**. Visual assessment of tumor growth in SKOV3 xenografts administered with two doses of $^{225}$Ac-Macropa- Isotype (0.6 μCi, 4 days apart)

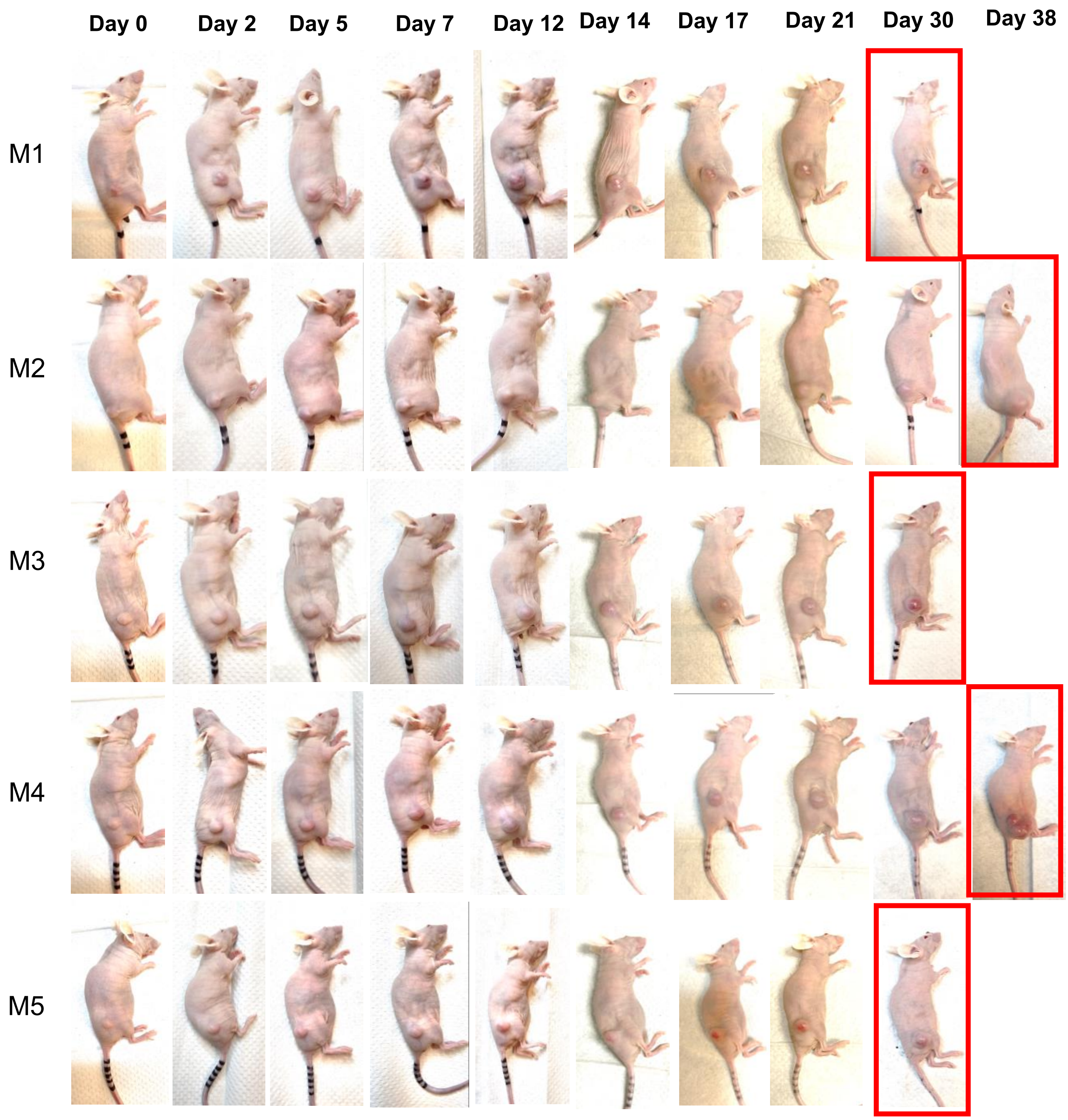


**S7. Tumor monitoring of Mirvetuximab Soravtansine**. Visual assessment of tumor growth in SKOV3 xenografts administered with two doses of Mirvetuximab Soravtansine (4.5 mg/Kg, 7 days apart)

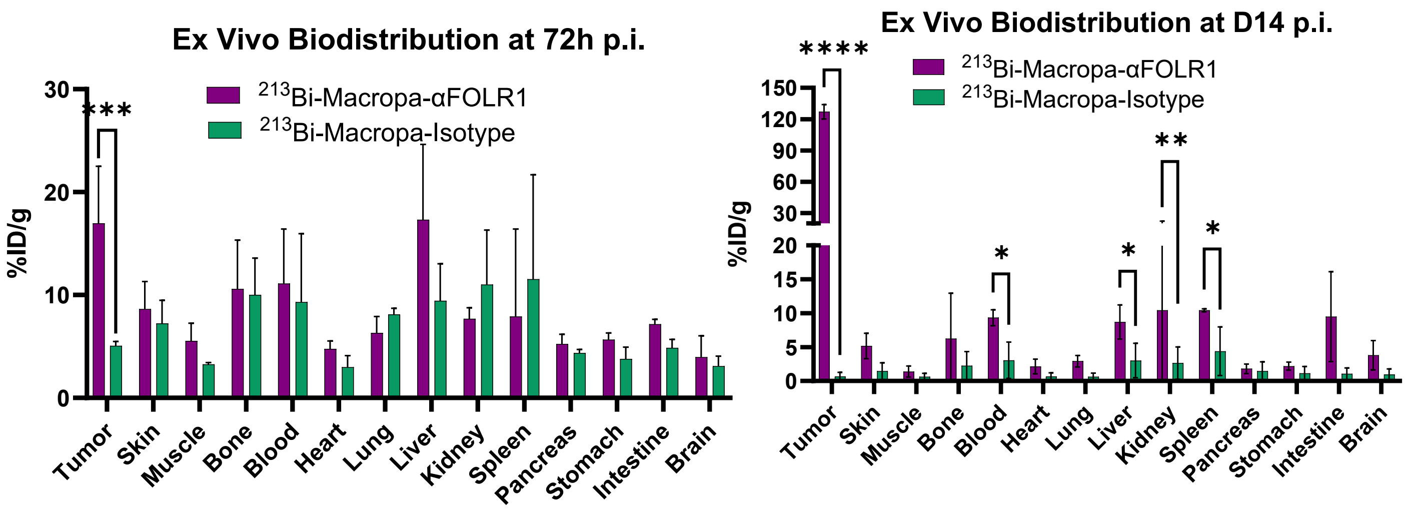


**S8. Ex vivo biodistribution of $^{213}$Bi-Macropa-αFOLR1 and $^{213}$Bi-Macropa-Isotype** Measurement of radioactivity the decay daughter of $^{225}$Ac, $^{213}$Bis, by gamma counting at (A) 72 h and (B) Day in SKOV3 mice receiving single 0.6 µCi and two doses (0.6 µCi), respectively**.**

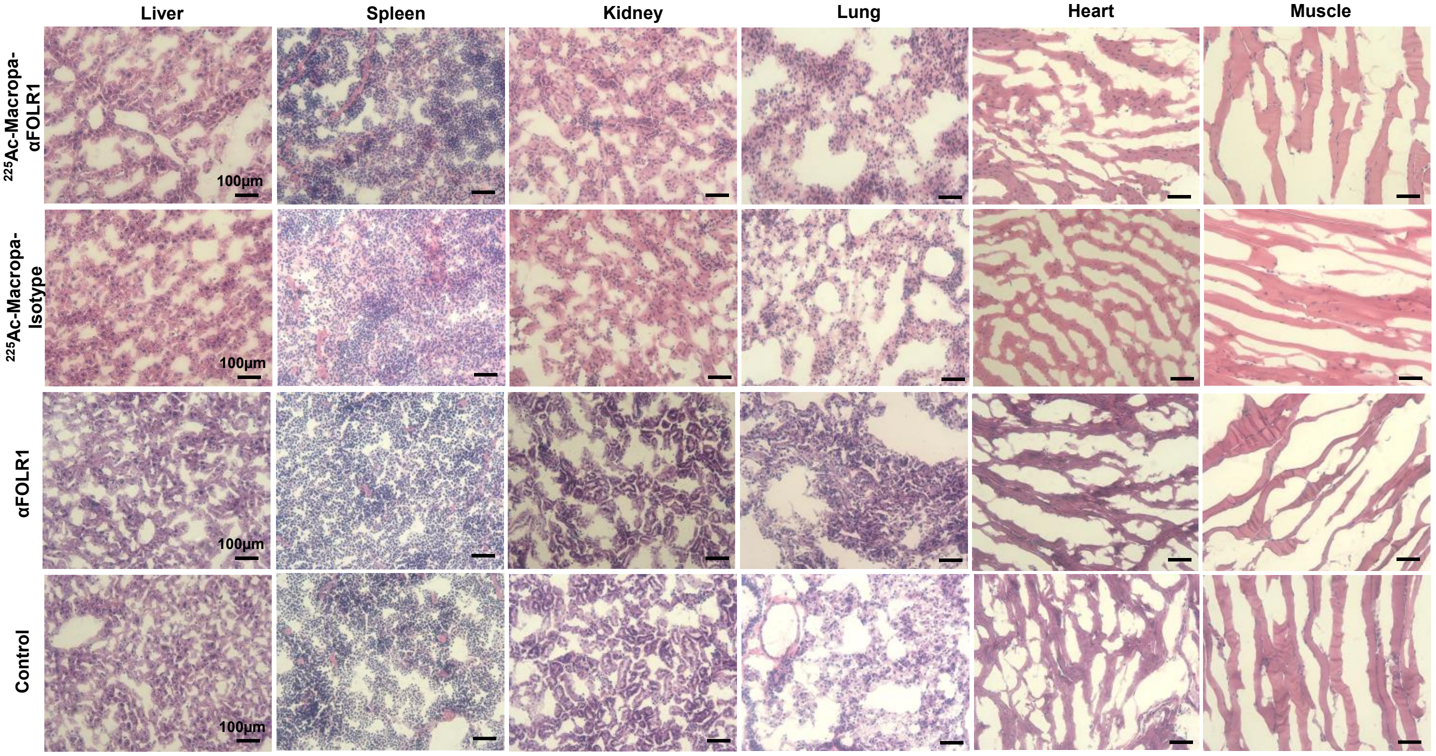


**S9. Histological evaluation of normal tissues by H&E on Day 14.** Histological evaluation of normal tissues at Day 14 by H&E staining for control, αFOLR1, $^{225}$Ac-Macropa-Isotype, $^{225}$Ac-Macropa-αFOLR1. (Images acquired at magnification 20X; Scale bar: 100µm)

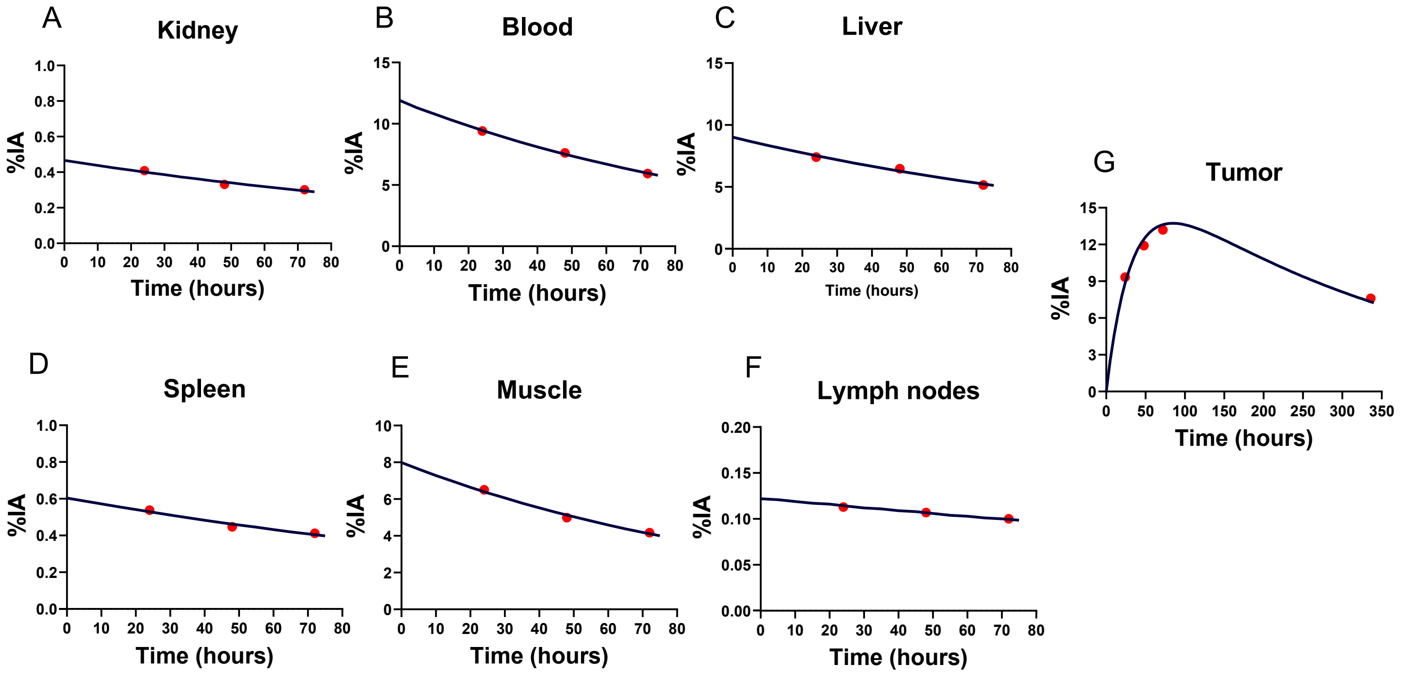


**S10. Time integrated activity curves for $^{225}$Ac-Macropa-αFOLR1.** TIACs generated by fitting percent average injected dose obtained from PET imaging data at 24. 48 and 72h in monoexponential function for kidney, blood, liver, spleen, muscle, and lymph nodes and biexponential fitting for tumors.

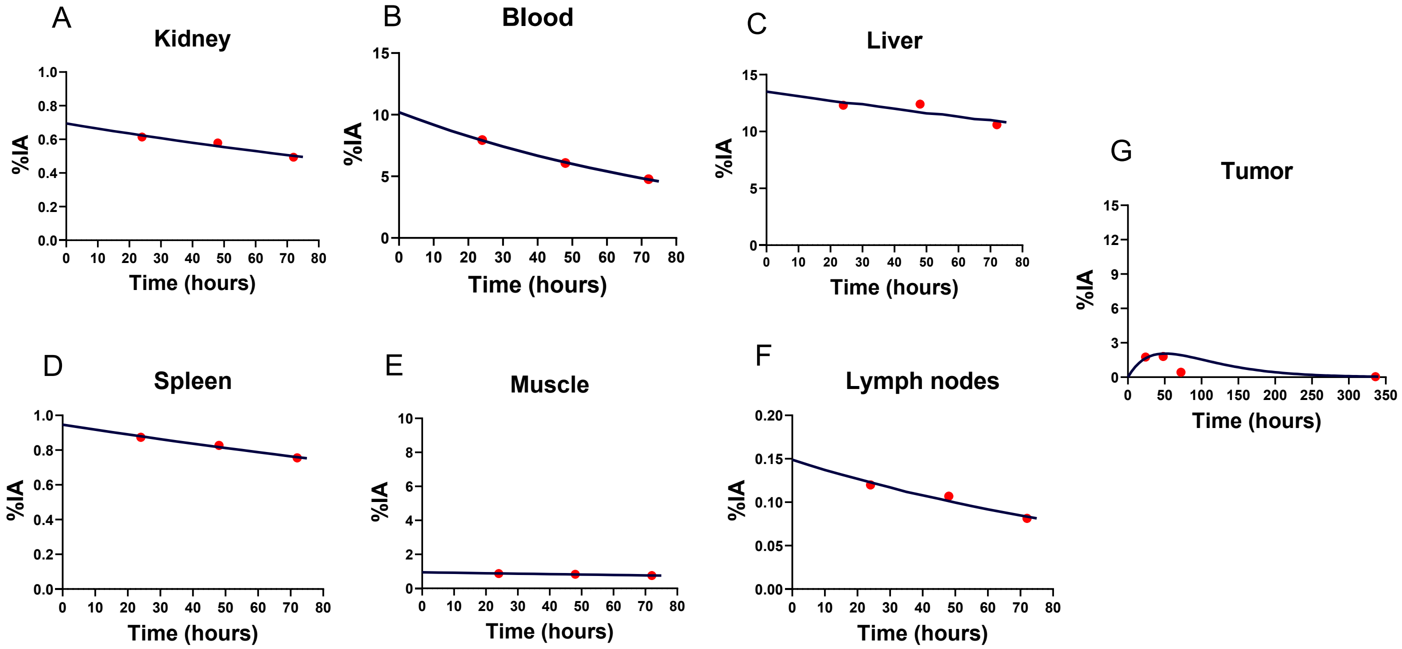


**S11. Time integrated activity curves for $^{225}$Ac-Macropa-Isotype.** TIACs generated by fitting percent average injected dose obtained from PET imaging data at 24, 48 and 72h in monoexponential function for kidney, blood, liver, spleen, muscle, and lymph nodes and biexponential fitting for tumors.

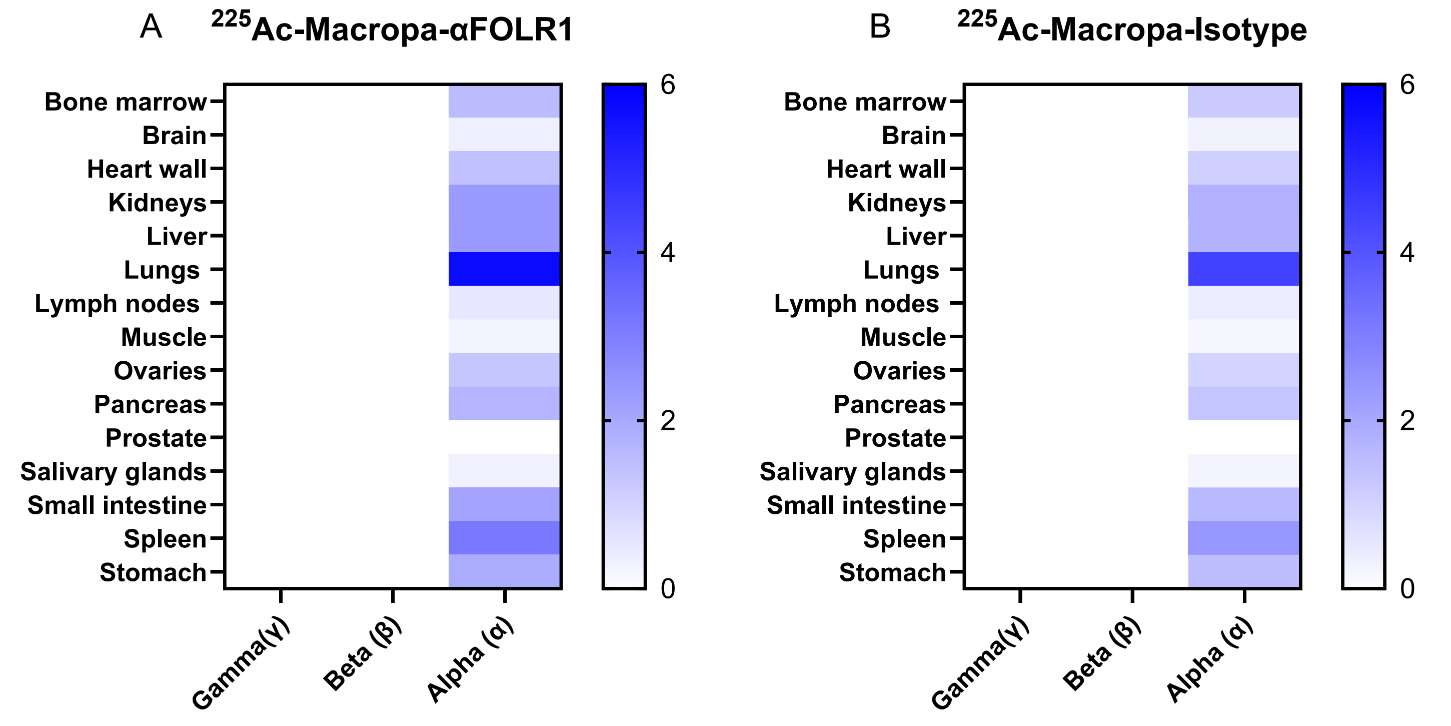


**S12. Cross absorbed dose for $^{225}$Ac-Macropa-αFOLR1 and $^{225}$Ac-Macropa-Isotype.** Heat map demonstrating the cross absorbed dose for alpha (α), beta (β), and gamma (γ) for different organs.

**Table 1.** Biokinetic fitting parameters for $^{225}$Ac-Macropa-αFOLR1

| $^{225}$Ac-Macropa-FOLR1 | $R^2$ | TIAC [h] | $A_0$(%) | $\lambda_1$($h^{-1}$) | $\lambda_2$($h^{-1}$) |
|---|---|---|---|---|---|
| **Kidneys** | 0.9503 | 0.7346 | 0.4664 | 0.0035 | |
| **Blood (classic ICRP)** | 0.9973 | 12.4152 | 11.9060 | 0.0067 | |
| **Liver** | 0.9799 | 11.9455 | 9.0177 | 0.0047 | |
| **Spleen** | 0.9512 | 1.0826 | 0.6045 | 0.0027 | |
| **Muscle** | 0.9863 | 8.6564 | 7.9881 | 0.0063 | |
| **Lymph nodes - systemic** | 0.9809 | 0.4240 | 0.1224 | 0.0000 | |
| **Tumor** | 0.9571 | 60.7333 | 19.3494 | 0.0000 | 0.0280 |

**Table 2**. Biokinetic fitting parameters for $^{225}$Ac-Macropa-Isotype

| $^{225}$Ac-Macropa-Isotype | $R^2$ | TIAC [h] | $A_0$(%) | $\lambda_1$($h^{-1}$) | $\lambda_2$($h^{-1}$) |
|---|---|---|---|---|---|
| **Kidneys** | 0.9378 | 1.5394 | 0.6946 | 0.0016 | |
| **Liver** | 0.7229 | 45.2410 | 13.5162 | 0.0001 | |
| **Blood (classic ICRP)** | 0.9993 | 9.6086 | 10.2057 | 0.0077 | |
| **Spleen** | 0.9817 | 3.1062 | 0.9460 | 0.0002 | |
| **Muscle** | 0.9323 | 9.1315 | 12.3356 | 0.0106 | |
| **Lymph nodes - systemic** | 0.9532 | 0.1850 | 0.1488 | 0.0052 | |
| **Tumor** | 0.9873 | 2.8529 | 96.3784 | 0.0162 | 0.0174 |

**Table 3.** Total absorbed dose per injected activity of $^{225}$Ac-Macropa- αFOLR1 and $^{225}$Ac-Macropa-Isotype

| | **$^{225}$Ac-Macropa-αFOLR1** | | **$^{225}$Ac-Macropa-Isotype** | |
|---|---|---|---|---|
| **Target Organ** | Absorbed Dose | Standard Deviation | Absorbed Dose | Standard Deviation |
| | [mGy / MBq] | | | |
| **Adipose tissue** | 0.167 | 0.0437 | 0.13 | 0.0318 |
| **Adrenals** | 1.66 | 0.434 | 1.3 | 0.316 |
| **Bone - endosteal cells** | 0.607 | 0.16 | 0.471 | 0.116 |
| **Bone marrow - red (active)** | 1.6 | 0.421 | 1.24 | 0.305 |
| **Brain** | 0.377 | 0.0996 | 0.292 | 0.0723 |
| **Breast tissue** | 0.31 | 0.0815 | 0.242 | 0.0592 |
| **Bronchial basal cells** | 1.69 | 0.446 | 1.31 | 0.324 |
| **Bronchial secretory cells** | 1.69 | 0.446 | 1.31 | 0.324 |
| **Bronchiolar secretory cells** | 1.69 | 0.446 | 1.32 | 0.324 |
| **Colon - ICRP133** | 2.07 | 0.547 | 1.61 | 0.397 |
| **Colon - left** | 2.07 | 0.547 | 1.61 | 0.397 |
| **Colon - rectosigmoid** | 2.07 | 0.547 | 1.6 | 0.397 |
| **Colon - right** | 2.07 | 0.547 | 1.61 | 0.397 |
| **Esophagus** | 1.97 | 0.518 | 1.53 | 0.376 |
| **ET1 airway basal cells** | 0.309 | 0.0815 | 0.239 | 0.0592 |
| **ET2 airway basal cells** | 0.309 | 0.0815 | 0.239 | 0.0592 |
| **Extrathoracic region - ICRP133** | 0.309 | 0.0815 | 0.239 | 0.0592 |
| **Eye lens** | 0.00128 | 0.000269 | 0.00112 | 0.000224 |
| **Gallbladder wall** | 0.322 | 0.0816 | 0.288 | 0.0709 |
| **Heart wall** | 1.46 | 0.385 | 1.14 | 0.279 |
| **Kidneys** | 9.4 | 2.79 | 16.6 | 7.99 |
| **Liver** | 24.9 | 7.24 | 87.2 | 72.2 |
| **Lung - ICRP133** | 5.79 | 1.53 | 4.48 | 1.11 |
| **Lungs (AI)** | 5.8 | 1.53 | 4.5 | 1.11 |
| **Lymph nodes - extrathoracic** | 0.56 | 0.148 | 0.434 | 0.108 |
| **Lymph nodes - systemic** | 12.1 | 10.4 | 5.47 | 1.52 |
| **Lymph nodes - thoracic** | 0.564 | 0.148 | 0.439 | 0.108 |
| **Lymphatic nodes - ICRP133** | 10.2 | 8.62 | 4.62 | 1.26 |
| **Muscle** | 1.9 | 0.457 | 1.93 | 0.389 |
| **Oral mucosa** | 0.309 | 0.0815 | 0.24 | 0.0592 |
| **Ovaries** | 1.34 | 0.354 | 1.04 | 0.257 |
| **Pancreas** | 1.77 | 0.465 | 1.38 | 0.337 |
| **Pituitary gland** | 0.309 | 0.0815 | 0.239 | 0.0592 |
| **Prostate** | 0 | 0 | 0 | 0 |
| **Salivary glands** | 0.309 | 0.0815 | 0.239 | 0.0592 |
| **Skin** | 0.525 | 0.139 | 0.407 | 0.101 |
| **Small intestine** | 2.13 | 0.563 | 1.65 | 0.409 |
| **Spleen** | 22.9 | 8.77 | 59 | 47.7 |
| **Stomach** | 1.97 | 0.518 | 1.54 | 0.376 |
| **Testes** | 0 | 0 | 0 | 0 |
| **Thymus** | 0.31 | 0.0815 | 0.241 | 0.0592 |
| **Thyroid** | 1.31 | 0.345 | 1.01 | 0.25 |
| **Tongue** | 0.309 | 0.0815 | 0.24 | 0.0592 |
| **Tonsils** | 0.309 | 0.0815 | 0.239 | 0.0592 |
| **Ureters** | 0.311 | 0.0815 | 0.244 | 0.0592 |
| **Urinary bladder wall** | 0.209 | 0.0549 | 0.162 | 0.0398 |
| **Uterus** | 0.309 | 0.0815 | 0.24 | 0.0592 |
| **Whole body target** | 2.05 | 0.58 | 4.44 | 3.06 |
| **Tumor** | 2430 | 362 | 139.33 | 14.4 |

**Table 4.** Distribution of total absorbed dose of $^{225}$Ac-Macropa- αFOLR1

| | $^{225}$**Ac-Macropa-αFOLR1** | | | | | |
|---|---|---|---|---|---|---|
| **Target Organ** | Gamma (self-dose) | Gamma (cross dose) | Beta (self-dose) | Beta (cross dose) | Alpha (self-dose) | Alpha (cross dose) |
| | | | | [mGy / MBq] | | |
| **Adipose tissue** | 0 | 0.00105 | 0 | 0.000706 | 0 | 0.165 |
| **Adrenals** | 0 | 0.0113 | 0 | 0.00694 | 0 | 1.64 |
| **Bone - endosteal cells** | 0 | 0.00289 | 0 | 0.00253 | 0 | 0.601 |
| **Bone marrow - red (active)** | 0 | 0.00594 | 0 | 0.00667 | 0 | 1.59 |
| **Brain** | 0 | 0.000547 | 0 | 0.00158 | 0 | 0.375 |
| **Breast tissue** | 0 | 0.00156 | 0 | 0.00129 | 0 | 0.307 |
| **Bronchial basal cells** | 0 | 0.0044 | 0 | 0.00703 | 0 | 1.68 |
| **Bronchial secretory cells** | 0 | 0.0044 | 0 | 0.00703 | 0 | 1.68 |
| **Bronchiolar secretory cells** | 0 | 0.00525 | 0 | 0.00705 | 0 | 1.68 |
| **Colon - ICRP133** | 0 | 0.00297 | 0 | 0.00866 | 0 | 2.06 |
| **Colon - left** | 0 | 0.00277 | 0 | 0.00866 | 0 | 2.06 |
| **Colon - rectosigmoid** | 0 | 0.00244 | 0 | 0.0087 | 0 | 2.06 |
| **Colon - right** | 0 | 0.00347 | 0 | 0.00864 | 0 | 2.06 |
| **Esophagus** | 0 | 0.00526 | 0 | 0.00821 | 0 | 1.95 |
| **ET1 airway basal cells** | 0 | 0.000369 | 0 | 0.00127 | 0 | 0.307 |
| **ET2 airway basal cells** | 0 | 0.00067 | 0 | 0.00131 | 0 | 0.307 |
| **Extrathoracic region - ICRP133** | 0 | 0.000658 | 0 | 0.00131 | 0 | 0.307 |
| **Eye lens** | 0 | 0.000278 | 0 | 0.000999 | 0 | 0 |
| **Gallbladder wall** | 0 | 0.0136 | 0 | 0.00178 | 0 | 0.307 |
| **Heart wall** | 0 | 0.00678 | 0 | 0.00657 | 0 | 1.45 |
| **Kidneys** | 0.00463 | 0.00664 | 0.0294 | 0.00995 | 6.99 | 2.36 |
| **Liver** | 0.02 | 0.00342 | 0.0942 | 0.0098 | 22.4 | 2.33 |
| **Lung - ICRP133** | 0 | 0.00729 | 0 | 0.0235 | 0 | 5.76 |
| **Lungs (AI)** | 0 | 0.00729 | 0 | 0.0235 | 0 | 5.77 |
| **Lymph nodes - extrathoracic** | 0 | 0.000951 | 0 | 0.000685 | 0 | 0.559 |
| **Lymph nodes - systemic** | 0.00395 | 0.00301 | 0.0478 | 0.00252 | 11.5 | 0.559 |
| **Lymph nodes - thoracic** | 0 | 0.00274 | 0 | 0.00235 | 0 | 0.559 |
| **Lymphatic nodes - ICRP133** | 0.0033 | 0.0028 | 0.0397 | 0.00235 | 9.55 | 0.559 |
| **Muscle** | 0.00136 | 0.000963 | 0.00688 | 0.00104 | 1.64 | 0.247 |
| **Oral mucosa** | 0 | 0.000955 | 0 | 0.00135 | 0 | 0.307 |
| **Ovaries** | 0 | 0.00272 | 0 | 0.00555 | 0 | 1.33 |
| **Pancreas** | 0 | 0.00747 | 0 | 0.00734 | 0 | 1.75 |
| **Pituitary gland** | 0 | 0.000394 | 0 | 0.00149 | 0 | 0.307 |
| **Prostate** | 0 | 0 | 0 | 0 | 0 | 0 |
| **Salivary glands** | 0 | 0.000637 | 0 | 0.00129 | 0 | 0.307 |
| **Skin** | 0 | 0.00106 | 0 | 0.00218 | 0 | 0.522 |
| **Small intestine** | 0 | 0.00401 | 0 | 0.00891 | 0 | 2.12 |
| **Spleen** | 0.0122 | 0.00493 | 0.0824 | 0.0132 | 19.6 | 3.15 |
| **Stomach** | 0 | 0.00758 | 0 | 0.00824 | 0 | 1.95 |
| **Testes** | 0 | 0 | 0 | 0 | 0 | 0 |
| **Thymus** | 0 | 0.0015 | 0 | 0.00132 | 0 | 0.307 |
| **Thyroid** | 0 | 0.00237 | 0 | 0.00541 | 0 | 1.3 |
| **Tongue** | 0 | 0.000933 | 0 | 0.0013 | 0 | 0.307 |
| **Tonsils** | 0 | 0.000703 | 0 | 0.00131 | 0 | 0.307 |
| **Ureters** | 0 | 0.00245 | 0 | 0.00131 | 0 | 0.307 |
| **Urinary bladder wall** | 0 | 0.00106 | 0 | 0.000877 | 0 | 0.207 |
| **Uterus** | 0 | 0.001 | 0 | 0.00129 | 0 | 0.307 |
| **Whole body target** | 0 | 0 | 0 | 0 | 0 | 0 |
| **Tumor** | 1.52 | | 10.16 | | 2420 | |

**Table 5.** Distribution of total absorbed dose of $^{225}$Ac-Macropa-Isotype

| | $^{225}$**Ac-Macropa-Isotype** | | | | | |
|---|---|---|---|---|---|---|
| **Target Organs** | Gamma (self-dose) | Gamma (cross dose) | Beta (self-dose) | Beta (Cross dose) | Alpha (self-dose) | Alpha (cross dose) |
| | | | [mGy / MBq] | | | |
| **Adipose tissue** | 0 | 0.00193 | 0 | 0.000551 | 0 | 0.128 |
| **Adrenals** | 0 | 0.0321 | 0 | 0.00563 | 0 | 1.27 |
| **Bone - endosteal cells** | 0 | 0.00392 | 0 | 0.00196 | 0 | 0.465 |
| **Bone marrow - red (active)** | 0 | 0.00813 | 0 | 0.00518 | 0 | 1.23 |
| **Brain** | 0 | 0.000482 | 0 | 0.00122 | 0 | 0.29 |
| **Breast tissue** | 0 | 0.00385 | 0 | 0.000999 | 0 | 0.238 |
| **Bronchial basal cells** | 0 | 0.00698 | 0 | 0.00544 | 0 | 1.3 |
| **Bronchial secretory cells** | 0 | 0.00698 | 0 | 0.00544 | 0 | 1.3 |
| **Bronchiolar secretory cells** | 0 | 0.0101 | 0 | 0.00551 | 0 | 1.3 |
| **Colon - ICRP133** | 0 | 0.00393 | 0 | 0.0067 | 0 | 1.6 |
| **Colon - left** | 0 | 0.00323 | 0 | 0.0067 | 0 | 1.6 |
| **Colon - rectosigmoid** | 0 | 0.00218 | 0 | 0.00673 | 0 | 1.6 |
| **Colon - right** | 0 | 0.00564 | 0 | 0.00669 | 0 | 1.6 |
| **Esophagus** | 0 | 0.00995 | 0 | 0.00639 | 0 | 1.51 |
| **ET1 airway basal cells** | 0 | 0.000458 | 0 | 0.000984 | 0 | 0.238 |
| **ET2 airway basal cells** | 0 | 0.000788 | 0 | 0.00101 | 0 | 0.238 |
| **Extrathoracic region - ICRP133** | 0 | 0.000774 | 0 | 0.00101 | 0 | 0.238 |
| **Eye lens** | 0 | 0.000347 | 0 | 0.000773 | 0 | 0 |
| **Gallbladder wall** | 0 | 0.0479 | 0 | 0.00276 | 0 | 0.238 |
| **Heart wall** | 0 | 0.0121 | 0 | 0.0051 | 0 | 1.12 |
| **Kidneys** | 0.0097 | 0.0159 | 0.0615 | 0.00774 | 14.6 | 1.83 |
| **Liver** | 0.0759 | 0.00323 | 0.357 | 0.00759 | 84.9 | 1.8 |
| **Lung - ICRP133** | 0 | 0.0117 | 0 | 0.0182 | 0 | 4.45 |
| **Lungs (AI)** | 0 | 0.0117 | 0 | 0.0183 | 0 | 4.47 |
| **Lymph nodes - extrathoracic** | 0 | 0.00115 | 0 | 0.000537 | 0 | 0.432 |
| **Lymph nodes - systemic** | 0.00172 | 0.00623 | 0.0209 | 0.00198 | 5.01 | 0.432 |
| **Lymph nodes - thoracic** | 0 | 0.00429 | 0 | 0.00182 | 0 | 0.432 |
| **Lymphatic nodes - ICRP133** | 0.00144 | 0.00564 | 0.0173 | 0.00184 | 4.17 | 0.432 |
| **Muscle** | 0.00143 | 0.00191 | 0.00726 | 0.000811 | 1.73 | 0.191 |
| **Oral mucosa** | 0 | 0.00105 | 0 | 0.00105 | 0 | 0.238 |
| **Ovaries** | 0 | 0.0024 | 0 | 0.0043 | 0 | 1.03 |
| **Pancreas** | 0 | 0.0189 | 0 | 0.0057 | 0 | 1.35 |
| **Pituitary gland** | 0 | 0.000414 | 0 | 0.00116 | 0 | 0.238 |
| **Prostate** | 0 | 0 | 0 | 0 | 0 | 0 |
| **Salivary glands** | 0 | 0.000747 | 0 | 0.001 | 0 | 0.238 |
| **Skin** | 0 | 0.00166 | 0 | 0.00169 | 0 | 0.404 |
| **Small intestine** | 0 | 0.00685 | 0 | 0.00691 | 0 | 1.64 |
| **Spleen** | 0.0351 | 0.00837 | 0.236 | 0.0102 | 56.3 | 2.44 |
| **Stomach** | 0 | 0.0191 | 0 | 0.00654 | 0 | 1.51 |
| **Testes** | 0 | 0 | 0 | 0 | 0 | 0 |
| **Thymus** | 0 | 0.00277 | 0 | 0.00102 | 0 | 0.238 |
| **Thyroid** | 0 | 0.00265 | 0 | 0.00419 | 0 | 1.01 |
| **Tongue** | 0 | 0.00102 | 0 | 0.00101 | 0 | 0.238 |
| **Tonsils** | 0 | 0.000819 | 0 | 0.00102 | 0 | 0.238 |
| **Ureters** | 0 | 0.00541 | 0 | 0.00102 | 0 | 0.238 |
| **Urinary bladder wall** | 0 | 0.00108 | 0 | 0.000679 | 0 | 0.16 |
| **Uterus** | 0 | 0.00104 | 0 | 0.001 | 0 | 0.238 |
| **Whole body target** | 0 | 0 | 0 | 0 | 0 | 0 |
| **Tumor** | 0.0813 | | 0.552 | | 131 | |
